# MLX balances metabolism and stress to suppress apoptosis and maintain spermatogenesis

**DOI:** 10.1101/2020.12.23.424063

**Authors:** Patrick A. Carroll, Pei Feng Cheng, Brian W Freie, Sivakanthan Kasinathan, Haiwei Gu, Theresa Hedrich, James A. Dowdle, Vivek Venkataramani, Vijay Ramani, Daniel Raftery, Jay Shendure, Donald E. Ayer, Charles H. Muller, Robert N. Eisenman

## Abstract

Male germ cell production is a metabolically-driven and apoptosis-prone process. Here we show that the glucose-sensing transcription factor MLX, and its binding partner MondoA, are both required for male fertility in the mouse, as well as survival of human tumor cells derived from the male germ line. Loss of *Mlx* results in altered metabolism as well as activation of multiple stress pathways and germ cell apoptosis in the testes. This is concomitant with dysregulation of the expression of male-specific germ cell transcripts and proteins, manifesting as oligoasthenoteratozoospermia (OAT). Our genomic and functional analyses identify loci directly bound by MLX involved in these processes, including metabolic targets, apoptotic effectors and obligate components of male-specific germ cell development. These *in vivo* and *in vitro* studies implicate MLX and other members of the proximal MYC network, such as MNT, in regulation of metabolism and differentiation, as well as in suppression of intrinsic and extrinsic death signaling pathways in both spermatogenesis and male germ cell tumors.

**Highlights:**

- The MAX-like bHLHLZ protein MLX is required for male fertility, but not embryonic development.
- MLX and its heterodimeric partner MondoA are each required for both male fertility and survival of male germ cell tumors.
- Genomic analysis identifies direct MLX targets associated with metabolism, stress and male germ cell development.
- Loss of MLX alters MYC network genome occupancy and transcriptional output.

## Introduction

The MYC/MAX/MXD network plays a critical role in both development and tumorigenesis, as major mediators of transcriptional regulation of growth, metabolism, proliferation, apoptosis and differentiation (for reviews see (Conacci-Sorrell et al., 2014);(Diolaiti et al., 2014; Kress et al., 2015)). This network is comprised of basic-helix-loop-helix-leucine zipper (bHLHLZ) transcription factors (TFs) generally associated with activation (MYC) or repression (MXD) that compete for an obligate heterodimerization partner (MAX) in order to bind DNA and influence expression of shared target genes(Carroll et al., 2018). Typically, MYC-MAX responds to mitogenic signals to activate Enhancer (E) box-containing promoters, whereas MXD-MAX responds to the loss of mitogenic signals or differentiation cues to repress the same targets. This allows the network to balance proliferative cues with cell cycle entry and exit.

The MAX-centered network exists within a larger network, containing MAX-Like protein X (MLX), MondoA (also known as MLX-interacting protein, MLXIP) and carbohydrate response element binding protein (ChREBP, also known as MondoB and MLXIPL) reviewed in (Billin and Ayer, 2006),(Havula and Hietakangas, 2012). MLX heterodimerizes with a subset of MXD proteins as well as the glucose-sensing MondoA and ChREBP but is unable to heterodimerize with either MAX or MYC. ChREBP-MLX (Iizuka et al., 2004) and MondoA-MLX heterodimers are major regulators of glucose-responsive transcription *in vitro* and *in vivo* ((Stoltzman et al., 2008) (Richards et al., 2018) (Ahn et al., 2016) (Ahn et al., 2019)) and MLX function has been linked to the response to metabolic stress in multiple organisms ((Havula et al., 2013) (Ma et al., 2005; Taniguchi et al., 2016) (Tamura et al., 2018)). Genetic ablation of MYC (Davis et al., 1993), MAX (Shen-Li et al., 2000) and MNT (Hurlin et al., 2003) (the most ubiquitously expressed MXD family member) results in embryonic or perinatal lethality (in the case of MNT), however loss of MondoA (Imamura et al., 2014) or ChREBP does not interfere with overt development (Iizuka et al., 2004). We previously demonstrated an obligate role for the MLX arm of the network in promoting survival of a wide-range of tumor cells with deregulated MYC by facilitating metabolic reprogramming and suppressing apoptosis. However, cells expressing endogenously regulated MYC were found to tolerate MondoA or MLX loss (Carroll et al., 2015).

Here we report the phenotype associated with loss of MLX during normal murine development. As with deletion of either *Mlxip* or *Mlxipl*, loss of *Mlx* is not detrimental to normal embryonic development or organismal viability. However, all male homozygous null animals (MLX^KO^) exhibit complete sterility with a dramatic increase in apoptosis of germ cells. We link this phenotype to a broad integrated transcriptional program mediated by MLX within the MYC network that facilitates metabolism and directly suppresses apoptosis.

## Results

### Homozygous deletion of *Mlx* is developmentally tolerated but results in male-only sterility

To examine potential developmental roles of *Mlx* in the mouse, we generated a targeting construct for deletion of exons 3-6 of murine *Mlx-α* encoding the bHLHLZ region required for dimerization and DNA binding (Figure 1A and Figure S1A). Upon constitutive heterozygous deletion of *Mlx*, we were able to obtain both *Mlx*^+/-^ males and females, indicative of developmental haplosufficiency, as reported for deletions of *Myc* or *Max*. However, upon mating of heterozygous males and females, we were surprised to discover that, unlike complete deletion of *Myc* or *Max* which results in embryonic lethality, homozygous loss of *Mlx* is well tolerated, resulting in offspring at the expected Mendelian frequencies (Figure 1B). *Mlx* null (MLX^KO^) mice of both sexes were indistinguishable from wild type (WT) mice, lived a normal lifespan, and exhibited normal behavior, including copulation (data not shown). Similar to *Mlx*^*+/-*^ (HET) mice of either sex, *Mlx*^*-/-*^ females were able to breed successfully, whereas all *Mlx*^*-/-*^ males were infertile (Figure 1C).

**Figure 1.**
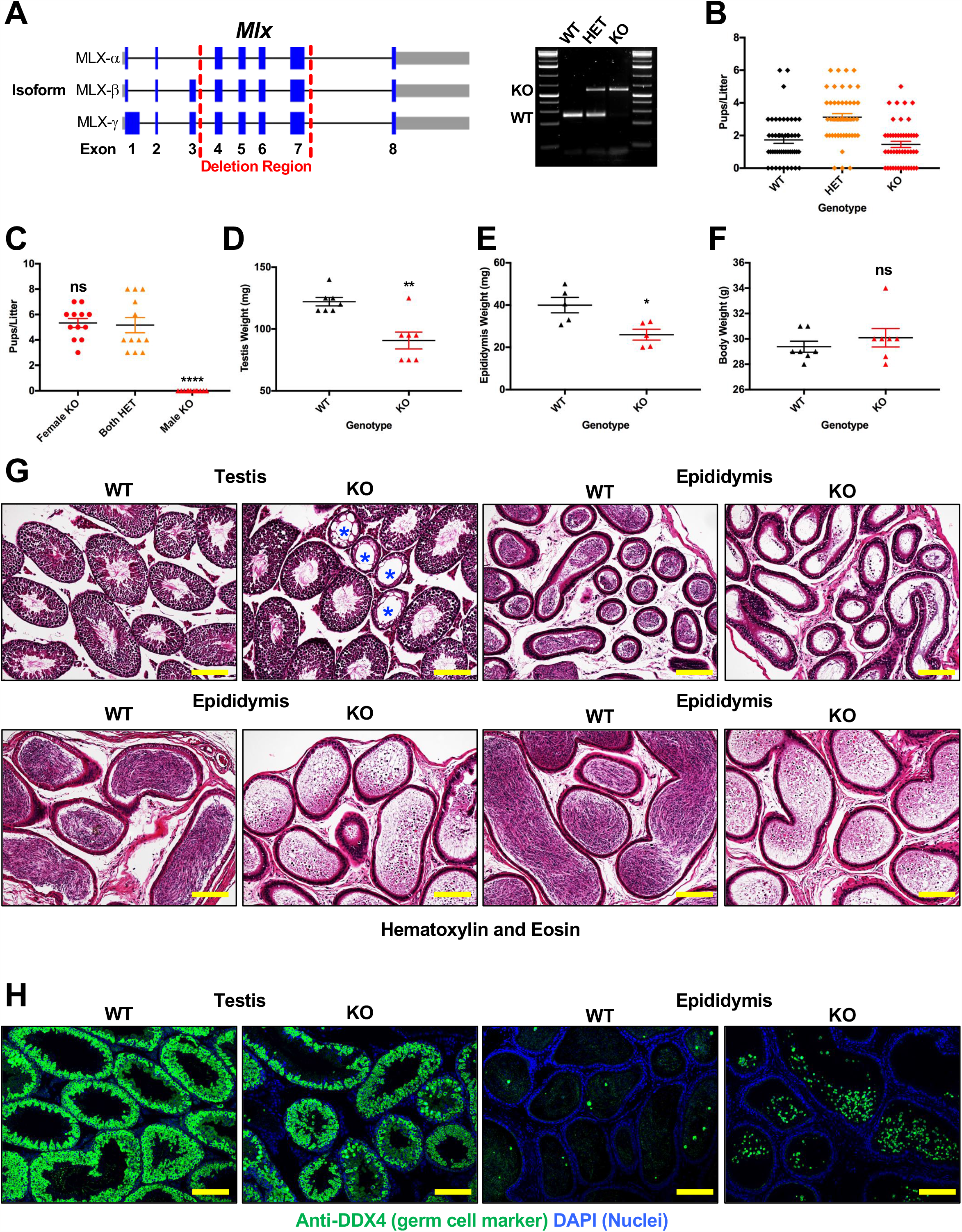
Cloning strategy and initial reproductive characterization of the MLX^KO^ mice. **A**. Cloning strategy for deletion of exons 3-6, encoding the DNA-binding domain, of murine *Mlx-α* with PCR products of the three potential genotypes WT, HET and KO shown to the right. **B**. Pups/litter of mating tests between MLX HET males and females (N=48). **C**. Mating test results from crossing the indicated sex and MLX genotype mice (N=12). **D**. Testis weight of WT versus MLX^KO^ mice (N=7). **E**. Epididymis weight of WT versus MLX^KO^ mice (N=5). **F**. Body weight of WT versus MLX^KO^ mice (N=7). **G**. Histological analysis of WT versus MLX^KO^ testis and epididymis stained with hematoxylin and eosin (100x, scale bar = 400uM). **H**. IF analysis of WT versus MLX^KO^ testis and epididymis stained for the indicated protein (100x, scale bar = 400uM). Shown for all is the Mean with SEM (* p<0.05, ** p<0.01, *** p<0.001, *** p<0.0001).

In accordance with the loss of fertility, MLX^KO^ testis and epididymis tissue were disrupted compared to WT mice. The weights of both testis and epididymis were significantly decreased in MLX^KO^ adult males (Figure 1D-E) despite normal body weight (Figure 1F). MLX^KO^ testes contained occasional abnormal, even acellular seminiferous tubules (marked with blue asterisks in Figure 1G), and MLX^KO^ epididymides contained decreased numbers of spermatozoa as well as populations of cells with an immature appearance compared to WT (Figure 1G). As shown in Figure 1H, the germ cell (GC) identity of these epididymal cells was confirmed by staining with the pan-GC cytoplasmic marker DDX4 (also known as VASA). These epididymal histological phenotypes of MLX^KO^ were not present in HET animals (Figure S1B) and were observed with varying severity from age P51 onward (Figure S1C-D).

### MLX^KO^ male mice are oligoasthenoteratozoospermic and sterility originates in the testes

To understand the basis for the testicular phenotype in the MLX^KO^ mice, we quantified aspects of the cellular biology of the testicular and epididymal GCs. Enumeration of daily sperm production rate (DSP) revealed both a significant decrease in DSP (Figure 2A), as well as diminished output of mature sperm to the epididymis in the MLX^KO^ compared with WT mice (Figure 2B), consistent with decreased production in the testis. The GCs that reached the epididymis in MLX^KO^ animals were confirmed to be immature as they exhibited both a lack of motility (Figure 2C) and abnormal morphology, as indicated by increased ratios of round vs. elongated spermatids (Figure 1H and 2D). These features match the clinical symptoms of oligoasthenoteratozoospermia (OAT) as defined by decreased sperm number, lack of motility and altered morphology of the sperm.

**Figure 2.**
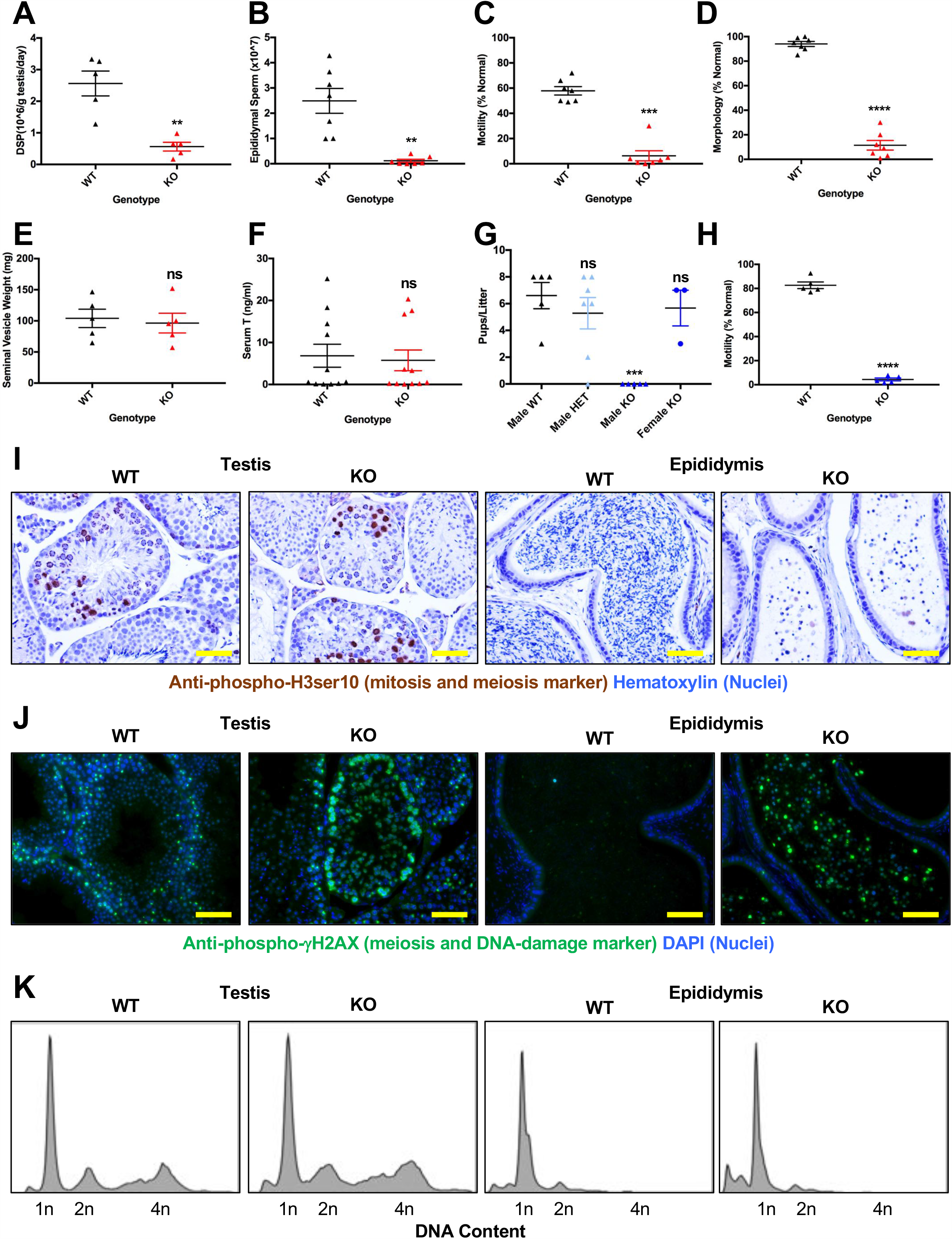
Fertility traits of WT versus MLX^KO^ and MondoA^KO^ mice. **A**. Daily Sperm Production (DSP) rate from WT versus MLX^KO^ mice (N=5). **B**. Epididymal sperm counts from WT versus MLX^KO^ mice (N=7). **C**. Percent normal motility of epididymal sperm from WT versus MLX^KO^ mice (N=7). **D**. Percent normal morphology of epididymal sperm from WT versus MLX^KO^ mice (N=7). **E**. Seminal vesicle weight from WT versus MLX^KO^ mice (N=5). **F**. Serum testosterone (T) from WT versus MLX^KO^ mice (N=11). **G**. Mating test results from crossing the indicated sex and MondoA genotype mice (N=3-6). **H**. Percent normal motility of epididymal sperm from WT versus MondoA^KO^ mice (N=5). **I**. IHC analysis of WT versus MLX^KO^ testis and epididymis stained for the indicated protein (200x, scale bar = 200uM). **J**. IF analysis of WT versus MLX^KO^ testis and epididymis stained for the indicated protein (200x, scale bar = 200uM). **K**. Flow cytometry analysis of single cell suspension from testis and epididymis of WT versus MLX^KO^ mice stained for DNA. Shown for all is the mean +/- SEM (* p<0.05, ** p<0.01, *** p<0.001, *** p<0.0001).

To determine whether a defect in androgen production correlated with the loss of fertility in MLX^KO^ males, we quantified serum testosterone (T) between WT and KO animals. Consistent with the lack of a change in the size of the seminal vesicle (Figure 2E), a testosterone responsive tissue, we did not detect a change in serum T levels between genotypes (Figure 2F). In contrast, alterations in levels of important metabolites were detected upon LC-MS/MS metabolomic analysis of serum from WT versus KO males. Partial least squares discriminant analysis (Figure S2A for PLS-DA), carried out using MetaboAnalyst (Xia et al., 2009), revealed multiple metabolites with a variable importance to projection (VIP) score of greater than 1. The top 20 are shown in Figure S2B and the Heat Map in Figure S2C. Ribose-5-phosphate, pyruvate, and lactate were increased in MLX^KO^ serum (Figure S2B-C), indicative of enhanced whole-body glycolysis. MLX^KO^ mice also exhibited alterations to metabolite levels associated with amino acid oxidation, such as decreased valine and a buildup of two downstream metabolites, 3-amino-isobutyrate and 2-hydroxy-isovaleric acid. Augmented glycolysis and alterations to oxidative substrates have also been reported in mice lacking MondoA, or treated with a chemical inhibitor of MondoA (Imamura et al., 2014) (Ahn et al., 2016)). MondoA-MLX heterodimers are known to act through their downstream target, TXNIP, which in turn suppresses glycolysis (Stoltzman et al., 2008). Taken together, these results indicate that deletion of *Mlx* leads to a change, not in the production of T, but in whole-body metabolism consistent with loss of MondoA-MLX activity, as well as alterations in proper testicular and epididymal tissue homeostasis.

As MLX functions as a transcription factor in concert with its heterodimeric-binding partners, MLX interacting proteins, we gauged the requirement of either *Mlxip* (encoding MondoA) or *Mlxipl* (encoding ChREBP) for male fertility. As shown in Figure 2G, deletion of the gene encoding MondoA, results in male-only sterility, while, consistent with previous publications (Iizuka et al., 2004), deletion of *Mlxipl* did not (data not shown). Interestingly, in contrast to the MLX^KO^ spermatozoa, which appeared immature, the spermatozoa from MondoA^KO^ mice appear normal (data not shown) and are produced at normal number (Figure S2D) but, are completely non-motile (asthenospermic) (Figure 2H). This supports a direct transcriptional requirement for MondoA-MLX activity in male fertility and also suggests that MLX has functions independent of MondoA in the context of spermatogenesis.

Given the phenotypic differences between the MondoA and the MLX deleted mice we sought to specifically determine the stage of the defect in spermatogenesis in the MLX^KO^ mice. Staining for phospho-Histone H3ser10 (to detect mitotic and meiotic cells) indicated that GCs from both WT and KO testes could undergo successful meiosis in the testis (Figure 2I). WT testis showed the expected stage-specific expression of phospho-*γ*H2AX (marker of meiosis and DNA damage) decreasing with differentiation and present only in rare epididymal GCs. In contrast, the MLX^KO^ tissue display disrupted expression and the shedding of immature, phospho-*γ*H2AX+ cells into the epididymis (Figure 2J). We also observed ectopic epididymal staining for the pan-GC marker DDX4 (Figure 1H), which is normally only detected at low levels due to the removal by phagocytosis of spermatid cytoplasts or residual bodies possessing this marker, during the transition from round to elongated spermatids.

As the cell markers cytoplasmic DDX4 and phospho-*γ*H2AX are normally lost upon completion of meiosis, we tested whether testicular and epididymal cells from MLX^KO^ mice were arresting in meiosis. All stages of meiosis (1, 2 and 4n DNA content) were observed in the testis and epididymis of both WT and MLX^KO^ mice (Figure 2K) indicating that the cells transiting to the epididymis of MLX^KO^ mice, although having significantly reduced total cell numbers compared to WT (Figure 2B), were post-meiotic (predominantly 1n). However, the MLX^KO^ epididymal population displayed increased sub-1n DNA content, indicative of DNA fragmentation (Figure 2K), consistent with phospho-*γ*H2AX expression. In conclusion, MLX^KO^ males exhibit OAT, with decreased testicular spermatid production accompanied by impaired transition from round to elongated spermatid morphology. These immature post-meiotic GCs maintain meiosis and stress markers and are shed to the epididymis where they appear to undergo apoptosis.

### Expression profiling reveals decreased spermatogenesis, altered metabolism and increased stress in MLX^KO^ testes

To gauge the altered transcriptome of MLX^KO^ tissue, we used RNA-Seq to profile the RNA of whole testes from age-matched littermates of WT verified fertile breeders versus MLX^KO^ males (n=3 pairs). As shown in Figure 3A, Principal Component Analysis (PCA) of these samples indicates that they group according to genotype. We identified 4688 differentially expressed genes (DEGs) upon loss of MLX (2282 up and 2406 down). (Figure 3B, and Table S1). Gene set enrichment analysis (GSEA) indicated enrichment for only the Spermatogenesis Hallmark Gene Set in the WT testes, which correlates with normal spermatogenesis in these mice (Figure 3C). However, compared with WT, the MLX^KO^ tissue is enriched for gene expression signatures related to multiple metabolic pathways including fatty acid metabolism, glycolysis, and oxidative phosphorylation (Figure 3C and Table S2). We also noted enrichment for stress pathways in the MLX^KO^ tissue, including inflammatory and interferon responses, TNF*α*, NF*κ*B, and JAK-STAT signaling, and apoptosis (Figure 3C and Supplemental Table 2).

**Figure 3.**
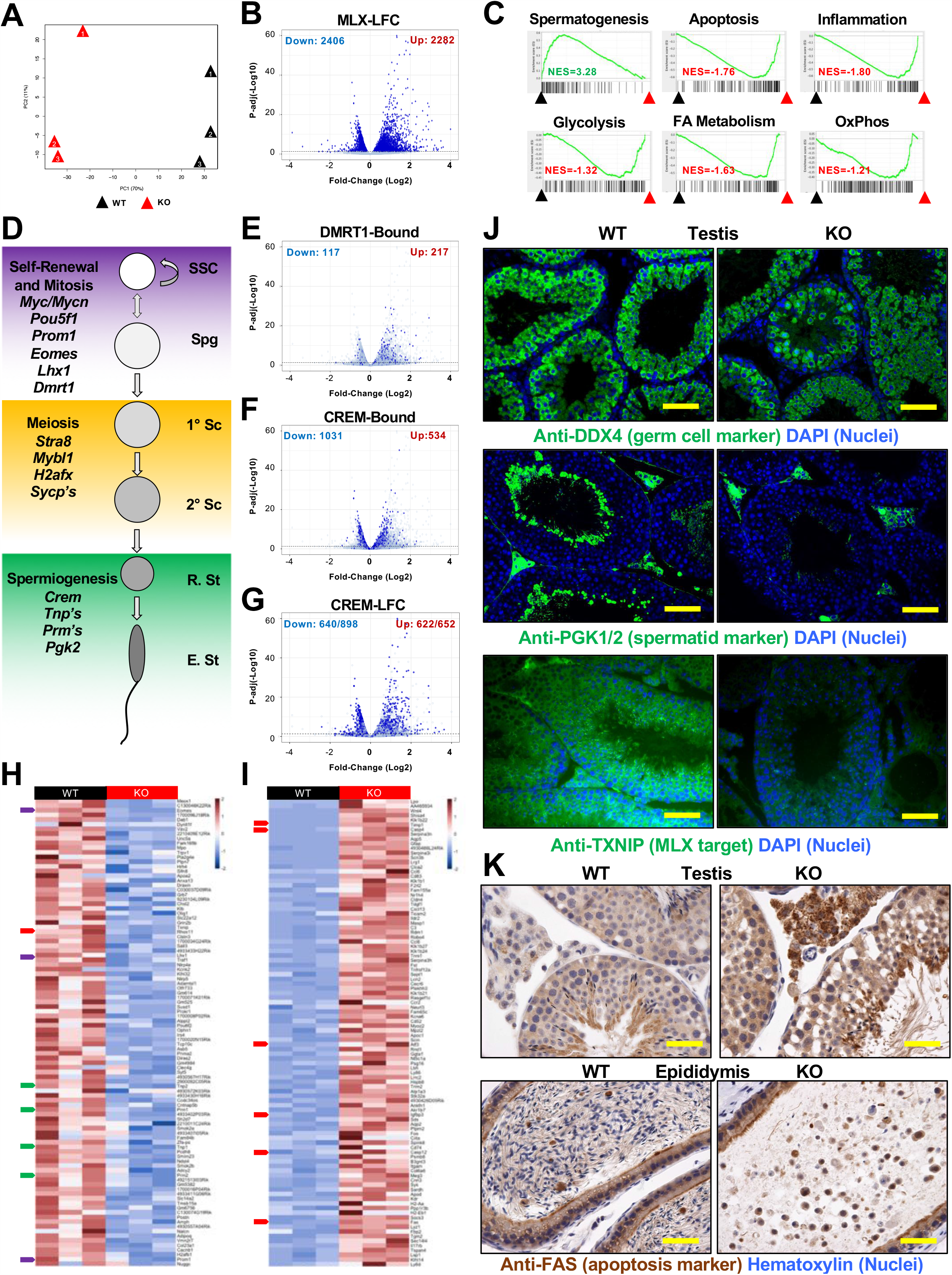
RNA profiling of testes from WT versus MLX^KO^ mice. **A**. Primary component analysis (PCA) of RNA-seq data from WT versus MLX^KO^ whole testes tissue. **B**. Volcano plot showing LogFC by –Log10 p-value of pair-wise analyzed RNA-seq data from WT versus MLX^KO^ whole testes tissue with significantly (p<=0.05) up/down differentially expressed genes (DEGs) indicated in dark blue. **C**. Gene set enrichment analysis (GSEA) of RNA-seq data with representative up/down categories shown (see supplemental Table 2 for complete list). **D**. Schematic of a simplified version of spermatogenesis broken into 3 major functions, Self-Renewal/Mitosis, Meiosis, and Spermiogenesis with the indicated stage-specific markers shown (Cell Type Abbreviations: SSC: Spermatagonial Stem Cell, Spg: Spermatagonia, Sc: Spermatocyte, R. St: Round Spermatid, E. St: Elongating Spermatid). **E**. Volcano plots showing LogFC by –Log10 p-value of pair-wise analyzed RNA-seq data (WT versus MLX^KO^) with significantly (p<=0.05) up/down genes indicated and blue marking those genes contained within the subgroups indicated from published datasets (DMRT1-Bound (Murphy et al., 2010)), CREM-Bound (Martianov et al., 2010)) and CREM-LFC (Kosir et al., 2012)). **F-G**. Heat Maps of top 100 down-regulated genes (**F**) in the MLX^KO^ testes and Heat map of top 100 up-regulated genes (**G**) in the KO testes. Colored arrows indicated DEGs associated with SSC/Spg function (Purple), Spermiogenesis (Green) and Apoptosis (Red). **H**. IF analysis of WT versus MLX^KO^ testis stained for the indicated proteins (200x, scale bar = 200uM). **I**. IHC analysis of WT versus MLX^KO^ testis and epididymis stained for the indicated protein (100x, scale bar = 400uM).

Spermatogenesis is a highly choreographed developmental program that can be separated into three broad categories enriched in three general cell types (spermatogonia (Spg), spermatocyte (Sc), and spermatid (St)), which undergo, respectively, self-renewal/mitosis, meiosis and spermiogenesis (see schematic in Figure 3D). These cellular states are lineage-specified through the activities of specific transcription factors and their targets, several of which are listed in figure 3D. Similar to GSEA, Enrichr analysis (Chen et al., 2013) (Kuleshov et al., 2016) for ChIP Set Enrichment (CHEA) was employed to identify transcription factors associated with differentially expressed genes from WT compared to MLX^KO^ testes. Most noteworthy is that genes downregulated in MLX^KO^ cells are significantly associated with loss of CREM and MYBL1 (required for spermiogenesis (Nantel et al., 1996) and meiosis (Bolcun-Filas et al., 2011)), respectively), while upregulated genes were associated with the more primitive spermatogonial transcription factors DMRT1 (Raymond et al., 2000)), OCT4 (Kehler et al., 2004), and MYC (Kanatsu-Shinohara et al., 2016)(Figure S3A). This suggests an incomplete block to proper spermatogenesis in MLX^KO^ germ cells with a loss of late markers and a buildup of more primitive markers of germ cell differentiation.

Volcano plots showing previously reported DMRT1-bound (Murphy et al., 2010) (Figure 3E) and CREM-bound (Martianov et al., 2010) (Figure 3F) targets from mouse testis overlapping with our RNA-Seq data are shown. DMRT1 is known to balance mitosis and meiosis-induction (Matson et al., 2010) and the trend towards upregulation of these targets (217 up versus 117 down) is consistent with the normal entry of these cells into meiosis, as evidenced by the presence of cells with 1N DNA content (see Figure 2K). The decreased expression of both MYBL1 and CREM targets is likely to be associated with stress during meiosis and spermiogenesis, respectively. Indeed, while the majority of CREM-bound targets are downregulated in our MLX^KO^ RNA-Seq analysis (1031 down versus 534 up), we find that a large fraction of differentially expressed genes in MLX^KO^ testes correspond to upregulated genes previously shown to be linked to deletion of *Crem* but not all of which are directly bound by CREM (622/652 upregulated genes, as opposed to 640/898 downregulated genes (Kosir et al., 2012) (CREM-LFC, Figure 3G). Our data implicate MLX in DMRT1- and CREM-regulated pathways critical for differentiation and survival of mammalian germ cells.

In order to highlight significantly altered transcripts from the GSEA/CHEA categories identified we generated heat maps for the top 100 downregulated and upregulated DEGs (Figure 3H-I). These include loss of both primitive SSC and Spg markers such as *Eomes, Lhx1* and *Prom1* (purple tabs), as well as the more differentiated spermatid markers *Tnp1, Tnp2, Prm1*, and *Prm2* (green tabs). We observe a concerted gain of apoptosis markers and effectors (red tabs), which include *Timp1, Casp4, Atf3, Igfbp3, Casp12 and Fas* upregulation (Figure 3I) as well as loss of *Txnip*, a pro-apoptotic protein that is MLX-dependent for expression (Stoltzman et al., 2008) (Figure 3H).

Immunoblotting of whole testes confirmed that a subset of these differentially expressed transcripts are also altered at the protein level. As shown in Figure S3B, upon MLX loss we observed decreased expression of both MLX dimerization partners MondoA and ChREBP, decreased expression of the known MondoA- or ChREBP-MLX target gene TXNIP, as well as decreased EOMES protein. However, consistent with GSEA enrichment for Apoptosis and Inflammation categories, we observed increased FAS (death receptor CD95) expression (Figure S3B) as well as reactivity with anti-mouse IgG HRP, indicative of resident immune cells, present in both WT and MLX^KO^, but increased in MLX^KO^ testes (Figure S3C).

We employed immunohistochemistry (IHC) on WT and MLX^KO^ testes and epididymides to ascertain *in situ* which populations of cells are altered upon loss of MLX. While both WT and MLX^KO^ testes stain positive for the pan-GC marker DDX4, the mature spermatid marker PGK2, as well as TXNIP, are significantly decreased in MLX^KO^ tissue (Figure 3J). The residual PGK signal detected in figure 3H is due to PGK1, present in interstitial and somatic cells. DDX4 expression confirms the GC fate of a large fraction of the testis cells. This, and the enrichment for DMRT1 targets among upregulated differentially expressed genes (Figure 3E), suggests that the proliferative Spg population is still present in MLX^KO^ testes. Consistent with this, the proliferation mark Ki67 is relatively unchanged in MLX^KO^ compared to WT tissue (Figure S3D). Taken together with the loss of markers such as PGK2, our data suggest that loss of MLX causes defects in the transition from spermatocyte to spermatid.

In contrast to the loss of PGK2, there is widespread staining for both the inflammation marker TIMP1 (Figure S3E) and FAS (Figure 3J and Figure S3F) in the seminiferous tubules and the interstitium of the testes, as well as the shed germ cells present in the epididymides of MLX^KO^ mice. A higher magnification image is presented for FAS staining in Figure 3K demonstrating that the round, immature spermatids that are both positive for DDX4 and phospho-*γ*H2AX are indeed also expressing FAS. This indicates that the loss of germ cells associated with MLX^KO^ tissue results from activation of the extrinsic cell death pathway via FAS death receptor signaling coincident with inhibition of differentiation.

### Gene expression in fractionated testes cell populations

We next fractionated the testicular tissue to remove interstitial (stromal and immune) cells from the seminiferous tubules (comprised predominantly of GCs as well as Sertoli cells) in order to assess protein expression. In comparison with WT, fractionated MLX^KO^ tubule cells show complete loss of MLX and strongly decreased expression of both MLX dimerization partners, MondoA and ChREBP. Surprisingly, these cells also exhibit moderately decreased expression of the immature spermatogonial stem cell (SSC) markers MYCN, MAX and OCT4, with no change in the expression of the MYC-antagonist MNT (Figure 4A). We had also noted diminished expression of the SSC marker EOMES in whole testes (Figure S3B). Importantly, siRNA against MLX resulted in similar changes in the male GC tumor (MGCT) cell line NTera2 (Figure 4B), supporting a cell-autonomous role for MLX in regulating the expression of these SSC markers. This suggests that MLX may impact stem cell function in MGCs and not just subsequent differentiation.

**Figure 4.**
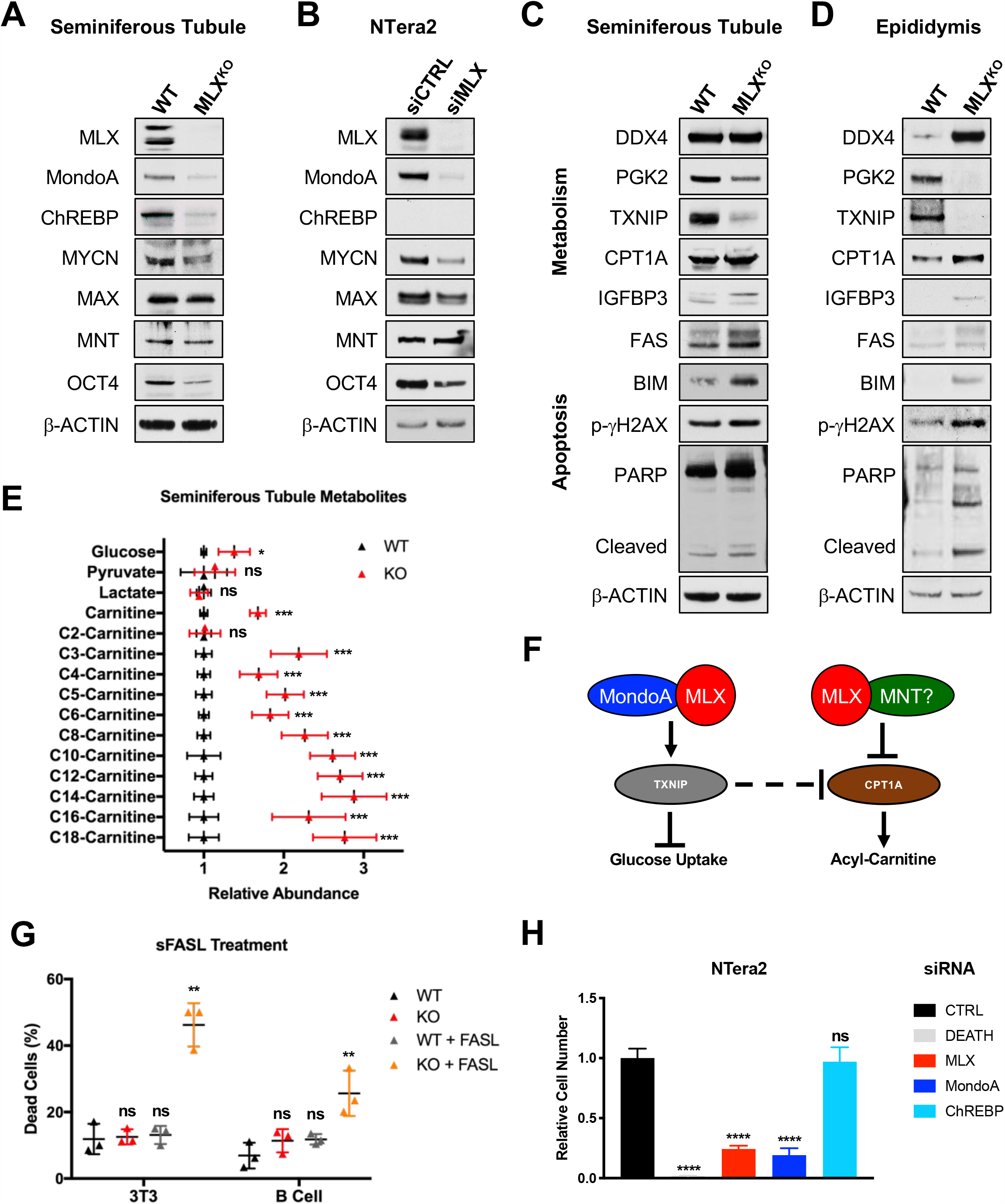
Molecular, biochemical and functional validation of GSEA categories from WT versus MLX^KO^ mice. **A-D**. Western blot analysis of isolated seminiferous tubule cells from WT versus MLX^KO^ mice. **B** NTera2 treated with siCTRL or siMLX. **C**. seminiferous tubule and **D**. epididymal cells from WT versus MLX^KO^ mice probed for the indicated proteins. **E**. Liquid Chromatography mass spectrometry (LC-MS) relative abundance data for the indicated metabolite from isolated seminiferous tubule cells from WT versus MLX^KO^ mice (shown is the mean +/- SD, N=3 paired littermates with N=4 technical replicates). **F**. A hypothetical model of regulation of metabolic targets by MLX. **G**. Soluble FASL (sFASL) treatment of the indicated cell lines (N=3 biological replicates, shown is the percent dead cells with mean +/- SD). **H**. Relative viable cell number of the NTera2 cells after siRNA transfection with the indicated siRNA with siDEATH included as a control for siRNA transfection efficacy (N=4 independent experiments, shown is the mean +/- SEM). (* p<0.05, ** p<0.01, *** p<0.001, *** p<0.0001).

The expression of metabolic and stress targets identified by RNA-seq, as well as markers of spermatogenesis, were assessed in the seminiferous tubules by western blot analysis of isolated cells from WT and MLX^KO^ mice. As shown in Figure 4C, the known MLX target TXNIP is decreased and the marker of fatty acid beta-oxidation CPT1A is increased along with stress related proteins including FAS, BIM, IGFBP3 and phospho-*γ*H2AX concomitant with PARP cleavage, all of which are consistent with increased apoptosis. We also confirmed decreased expression of the mature germ cell marker PGK2, while the pan-GC marker DDX4 is not significantly altered. This further confirms a disruption of normal differentiation associated with elevated stress.

In contrast with our observations in seminiferous tubules, cells isolated from epididymides did not robustly express network or stem cell markers (data not shown). However, as shown in Figure 4D, MLX^KO^ epididymal cells did exhibit alterations to the same metabolic targets TXNIP and CPT1A, as well as elevated stress markers FAS, BIM, IGFBP3, phospho-*γ*H2AX and PARP cleavage. Epididymal cells from MLX^KO^ mice also maintained the immature germ cell marker DDX4 (which is normally absent from epididymal cells of WT mice) and, consistent with spermiogenic defects, they also lacked the mature marker PGK2. Interestingly, MLX appears to regulate many of the same proteins in cells isolated from the interstitium of the testes, supportive of a broad role for MLX in both stabilizing binding partners and regulating metabolic targets (Figure S4A).

### MLX regulates glucose and lipid metabolism and suppresses apoptosis

In order to gauge the functional consequences of alterations to metabolism associated with loss of MLX, targeted LC-MS/MS was utilized to monitor glycolytic and beta-oxidation metabolites in isolated seminiferous tubule cells. As shown in Figure 4E, we detected increased intracellular glucose, consistent with the diminished expression of TXNIP which is known to suppress glucose uptake, while there was no significant change in pyruvate or lactate levels, perhaps due to decreased expression of glycolytic enzymes that are targets of CREM (e.g. LDHA and LDHC). We also detected a significant increase in a number of acyl-carnitine species (the product of CPT1A enzymatic activity) (Figure 4E), while there was no change in acetyl-carnitine (C2-carnitine), consistent with diminished expression of CRAT (another CREM target down in the MLX^KO^ testes) as opposed to the general upregulation of fatty acid gene set in general, many of which are regulated by CREM (e.g. XBP1, SREBF1). These changes are consistent with a role for MLX as a transcriptional regulator of metabolism in the seminiferous epithelium predominated by germ cells. A hypothetical model for putative targets of MLX responsible for these changes is shown in Figure 4F, including the published positive correlation between MondoA-MLX and TXNIP (Stoltzman et al., 2008) and the inverse correlation between TXNIP and CPT1A (Chen et al., 2016).

To directly assess whether MLX plays a cell-autonomous role in regulating the FAS death receptor, we determined the effect of soluble FAS ligand (sFASL) on immortalized 3T3 cells and primary B cells derived from WT versus MLX^KO^ mice. As shown in Figure 4G, FASL selectively kills both MLX^KO^ 3T3 cells and primary B cells while minimally affecting WT cells under normal culture conditions. We note that the MLX^KO^ and WT 3T3 cells are equally viable under standard culture conditions. However, as we previously reported, MLX^KO^ 3T3 cells undergo rapid apoptosis following enforced MYC expression (Carroll et al., 2015). Importantly, the expression of FAS protein is also elevated in MLX^KO^ 3T3 cells and is suppressed by the reintroduction of any one of the three isoforms of MLX into null cells (Figure S4B). Reintroduction of MLX also stabilizes MondoA and MYC-MAX protein levels. Consistent with the established role of MondoA-MLX (Carroll et al., 2015), the MLX targets TXNIP, TOMM20, FASN are lost in the MLX^KO^ cells, but robustly re-established with reconstitution of MLX (Figure S4B). This indicates a direct role for MLX in both activation of metabolic targets and suppression of FAS levels and suggests that MLX loss sensitizes cells to context-dependent death, not only as a consequence of MYC activation, but also in response to environmental factors such as FASL and glucose levels. We surmise that MLX normally attenuates stress and apoptosis during spermatogenesis, a process involving high metabolic demand, dependent upon a glycolytic program driven by both MYC and MYCN (Kanatsu-Shinohara et al., 2016)), as well as directly modified by FASL-FAS signaling (Lee et al., 1997).

### Male germ cell tumors require MLX for survival

To extend our observations on the requirement for MLX in spermatogenesis and cellular survival, we silenced MLX, and its dimerization partners MondoA or ChREBP, in a MGCT cell line (NTera2). Knockdown of MLX or MondoA in NTera2 cells resulted in a significant reduction in both the expression of SSC markers (Figure 4B and Figure S4E) and in viability (Figure 4H), while siChREBP has no effect on NTera2 cells, as expected, since it is not expressed. This siRNA, along with those against MondoA or MLX successfully killed HepG2 cells, a ChREBP-dependent tumor cell line (Tong et al., 2009) (Figure S4C), and also led to loss of both MondoA and ChREBP target gene expression (Figure S4D). These data are consistent with both cell-type specific effects of MondoA versus ChREBP, as well as a cell-autonomous requirement for MondoA-MLX in germ cell tumors.

As shown by immunoblot, both NTera2 and HepG2 cells treated with siMLX exhibited loss of MLX-dependent metabolic targets, (Figure S4D-E) overlapping with the targets in the 3T3 cells (Figure S4B). Also similar to GCs *in vivo* (Figure 4A), the expression of MYCN and MAX decreased upon knockdown of MondoA or MLX in the NTera2 cells (Figure S4E). A broader analysis of MGCTs based on TCGA data shows that *MLX* correlates with *POU5F1*, encoding OCT4, as well as *MYCN* (Figure S4F). Moreover, an independent dataset from oncomine.org (Korkola et al., 2005) also indicates over-expression of *MLX, POU5F1, MYCN, MLXIP* (encoding MondoA), *MAX*, and *MYC* in MGCT (Figure S4G). IHC analysis of xenografts derived from NTera2 cells (Venkataramani et al., 2012) (Figure S4H), shows all of these factors co-expressed in the pluripotent and proliferative (Ki67+) region of the tumors. This supports a role for MondoA-MLX in viability of germ cell-derived tumor cells, suggesting that they coordinately regulate proliferation and stemness and suppress cell death.

### MLX directly regulates male germ cell development in coordination with MAX

To identify genomic binding sites for MLX and MAX, we carried out ChIP-Seq on WT and MLX^KO^ testes. MLX binding sites were associated with 855 individual gene loci, while MAX peaks were associated with 874 gene loci in the WT tissue and 627 gene loci in the MLX^KO^ tissue (association defined as within 5Kb of the TSS of a locus) (Figure 5A). Differentially expressed transcripts in WT versus MLX^KO^ testes, as detected by RNA-Seq, showed significant correlation with MLX and MAX binding (Figure 5B-C), suggesting that these factors contribute directly to the regulation of genes whose expression is altered upon MLX loss. While genes occupied by MAX or MLX in general tend to display decreased expression in MLX deleted testes, many MAX or MLX bound genes were also found to be upregulated subsequent to MLX loss (Figure 5D). A subset of gene promoters exhibited binding by both MAX and MLX (Figure 5A and Figure 5E). Among these, as shown by the tracks in Figure 5E, are promoters of previously reported targets such as *Txnip*, as well as loci encoding protamine and transition proteins (*Prm* and *Tnp*, respectively), key genes involved in spermatogenesis.

**Figure 5:**
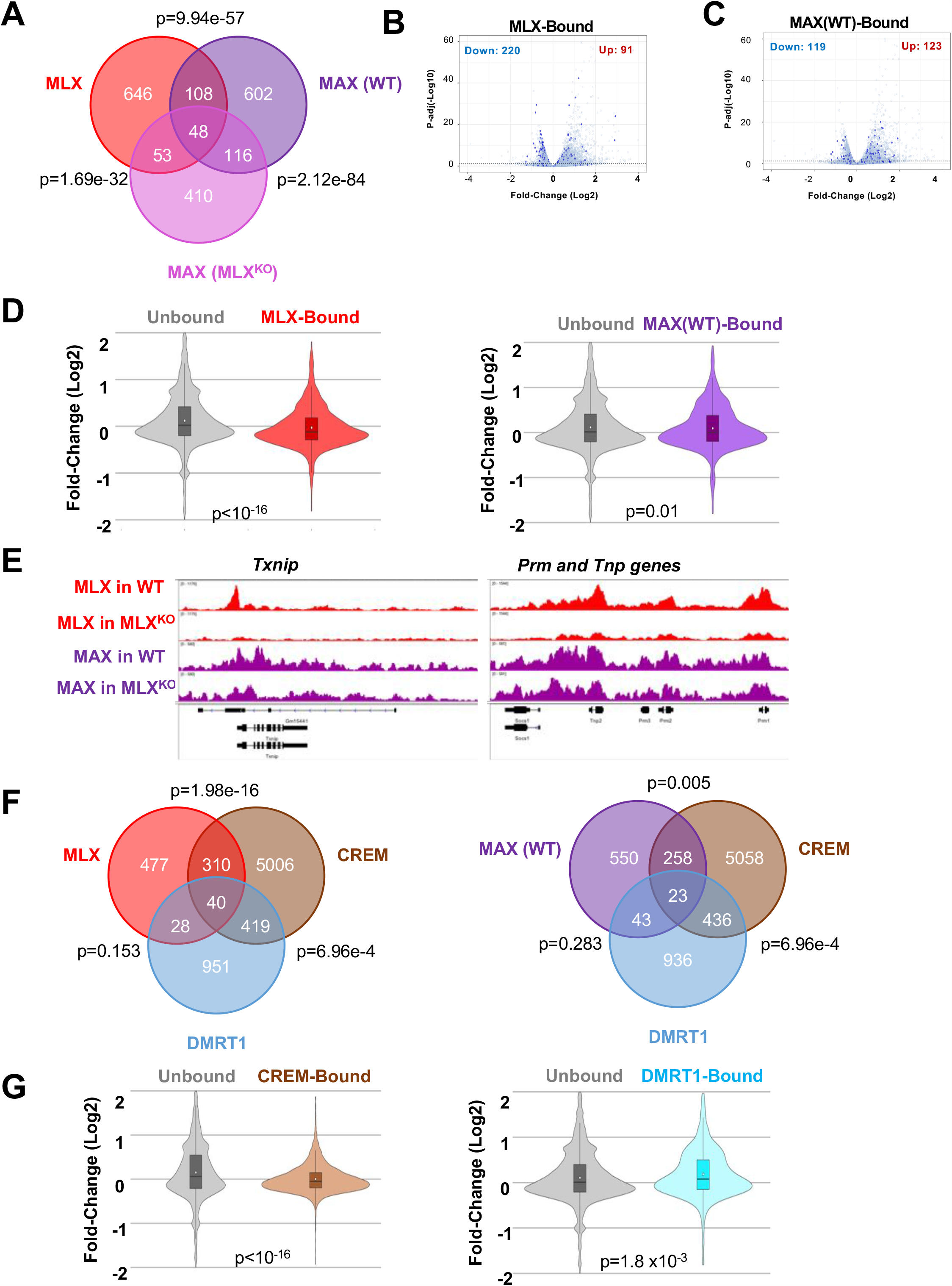
MLX directly regulates male germ cell (MGC-)specific targets. **A**. Venn Diagram of genes bound by MLX and MAX in WT and MLX^KO^ testes identified by ChIP-Seq (Hypergeometric test p-values shown) **B**. Volcano plot of MLX-bound loci (dark blue dots) from WT testis overlapping with RNA-Seq data (KS test p=0.000000e+00). **C**. Volcano plot of MAX-bound loci (dark blue dots) from WT testis overlapping with RNA-Seq data (KS test p=3.853399e-02). **D**. Violin plots comparing LFC of WT versus MLX^KO^ testes for MLX-bound vs unbound loci (left) and MAX-bound vs unbound loci (right) loci. (T-test). **E**. ChIP-Seq IGV tracks for *Txnip* and male germ cell-specific genes (*Prm* and *Tnp*) targets and TXNIP showing occupancy by MLX and MAX in WT and MLX^KO^ testes. **F**. Venn diagrams showing binding overlap of MLX (left) and MAX (right) with known CREM and DMRT1 target genes in testes. **G**. Violin plots comparing LFC upon MLX deletion for CREM-bound vs unbound (left) and DMRT-bound vs unbound (right) loci. (T-test).

Moreover, we find that MLX binding occurs at a group of genes previously reported to be targets of the transcriptional regulator CREM (Figure 5F). MAX binding is also apparent at CREM targets (Figure 5F). These targets display altered expression in MLX null testes relative to WT testes, as shown above (Figure 3E-G) and in Figure 5G. While the CREM transcription factor binds to a significant number of DMRT1 target genes, few DMRT1 regulated genes are also bound by MLX or MAX (Figure 5F). The CREM and DMRT1 transcription factors are considered to be essential mediators of spermatogenesis and spermiogenesis and the overlap between MLX/CREM binding is consistent with the observation that MLX loss phenocopies CREM loss of function and results in a similar pro-apoptotic phenotype in male germ cells (Nantel et al., 1996). We note that the testes-specific binding to key male germ cell specific genes (e.g. *Prm* and *Tnp*, respectively) (Figure 5E) is consistent with the observed male-specific fertility phenotype of MLX null mice, as female germ cells do not express protamines or transition proteins, associated with sperm-specific genomic compaction.

We also note that in testes harboring an *Mlx* deleted locus, MAX occupancy is altered in that MAX occupies only 19% (164/874) of its targeted gene loci detected in WT testes. In MLX^KO^ testes, MAX is present at 53 former MLX sites and occupies 410 new sites not occupied by either MLX or MAX in the WT testes (Figure 5A). These data suggest MAX and MLX complexes mutually influence their genomic binding sites or that loss of MLX results in changes in the representation of testes cell populations that possess distinct MLX and MAX binding signatures.

### MLX regulates metabolism and stress in multiple cell types

MLX, MAX and their major heterodimerization partners are expressed in multiple cell types in addition to testes. In order to assess common aspects of their binding and functions across tissues we surveyed their binding in primary B220+ splenic B cells stimulated with LPS from WT and MLX^KO^ mice. In contrast with the whole testis analysis, these samples have the advantage of comprising relatively homogeneous cell populations and were sensitive to FASL-induced killing (Figure 4G). In LPS stimulated WT B cells both MLX and MAX are both present at promoters with a strong propensity for association with canonical E-boxes (Figure S5A-B). Co-occupancy by MLX and MAX is exemplified by the tracks shown for the metabolic regulator TXNIP and the pro-apoptotic target, BIM (encoded by *Bcl2l11*) in B cells (Figure S5C). Changes in protein levels in WT vs. MLX^KO^ B cells recapitulate critical features observed in MLX^KO^ germ cells, such as loss of TXNIP, destabilization of MYC, induction of stress indicators such as BIM and markers of DNA damage and apoptosis (Figure S5D). Moreover, as observed in testes, MAX occupancy shifts between WT and MLX^KO^ cells showing significant binding to sites not previously occupied in MLX WT cells (Figure S5E).

### Both MLX and MNT bind metabolic and stress targets shared with MAX

Apparent alterations in genomic occupancy by MAX in MLX^KO^ cells compared to WT cells could reflect a shift in heterodimer formation by binding partners untethered from MLX, and therefore available to dimerize with MAX. To examine occupancy by other members of the MYC network we extended our ChIP-Seq analysis in WT and MLX^KO^ 3T3 cells to include the transcriptional repressor MNT which has been shown to independently dimerize with MAX as well as with MLX and also to form MNT-MNT homodimers (Lafita-Navarro et al., 2020)). MLX, MAX and MNT exhibit a similar genomic distribution of occupancy proximal and downstream of the TSS within regions significantly enriched for the E-Box sequence motif (Figure 6A-B). MAX binds to the largest share of loci while MLX and MNT occupy a subset of loci occupied by MAX (Figure 6C). Moreover, data from individual tracks indicate that multiple MYC network members can occupy the same promoter regions and exhibit subtle changes in occupancy upon MLX deletion (Figure 6D). Interestingly, as observed in both the testes and primary B cells (Figure 5A, Figure S5E), the genes bound by both MAX and MNT shift between WT and MLX^KO^ 3T3 cells (Figure S6A). Both MLX deletion and MNT deletion independently induce increased protein expression of BIM and the intrinsic stress response protein ATF4 (Figure 6E). This raises the possibility that MNT-MLX heterodimers contribute to the response to stress. As expected, deletion of MLX reduces levels of the well-established MondoA-MLX target TXNIP, however loss of MNT induces TXNIP suggesting that MNT may normally function with MAX or MLX to suppress TXNIP expression (Hypothetical Model in Figure 6F). Supportive of MNT-MLX crosstalk, we observe co-occupancy of MNT and MAX with MLX targets (*Arrdc4, Fasn* and *Atf4*) in both 3T3 cells (Figure S6B), as well as annotated datasets on ENCODE from the MLX-dependent (Figure S4C) human HepG2 cells (Figure S6C) showing binding for both *TXNIP* and *BCL2L11*, as well as *FASN*.

**Figure 6:**
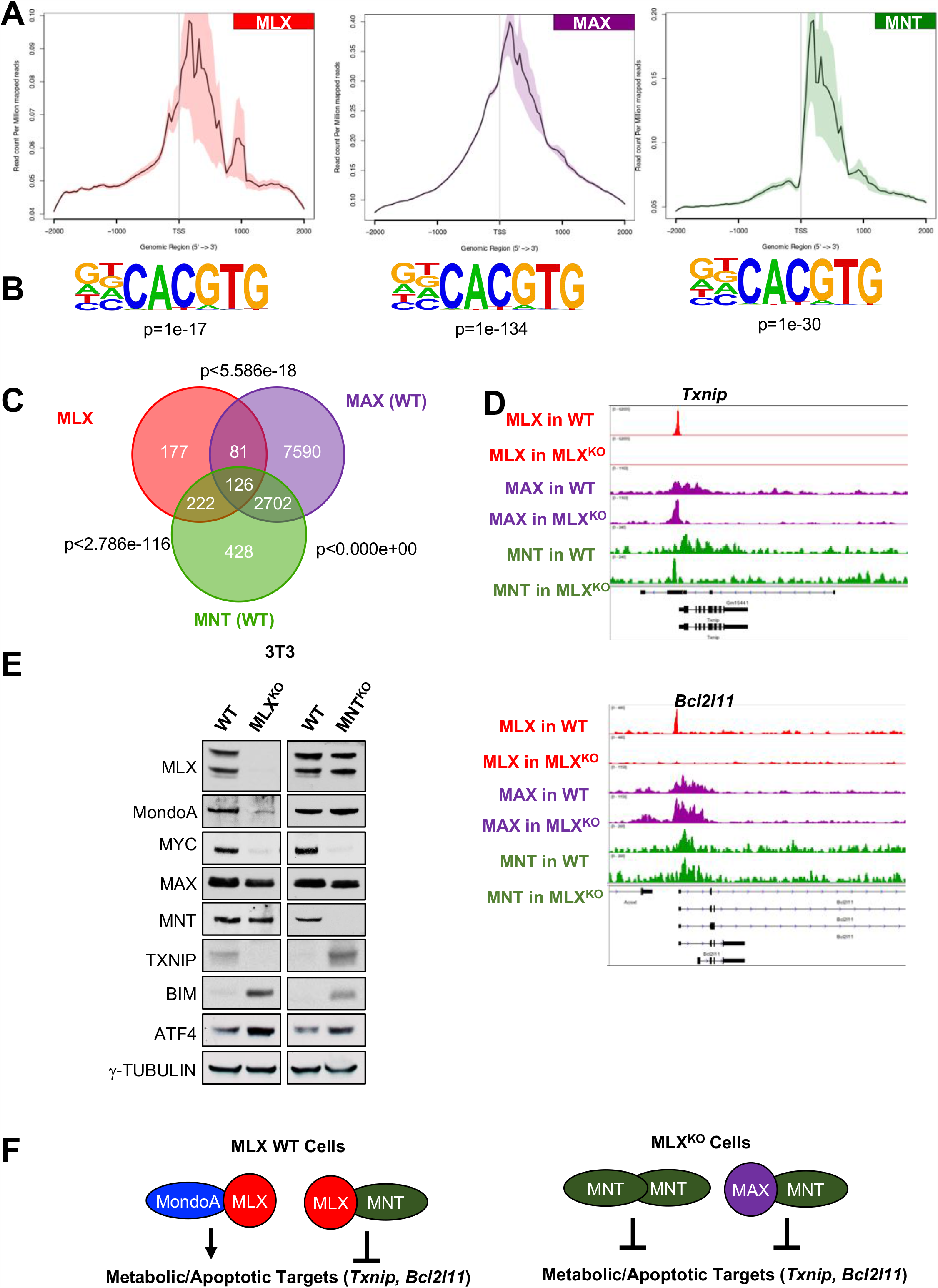
MLX shares numerous transcriptional targets with MAX and MNT. **A**. NGS plots for MLX, MAX and MNT in WT 3T3 cells. **B**. E-Box Motif from HOMER for MLX, MAX and MNT with the indicated p-value. **C**. Venn Diagram of the overlap of genes bound by MLX, MAX and MNT in 3T3 cells with p-value calculated from a hypergeometric test. **D**. IGV tracks for MLX, MAX and MNT on the *Txnip* and *Bcl2l11 (Bim)* promoters from WT and MLX^KO^ 3T3 cells. **E**. Western blots from WT versus MLX^KO^ and WT versus MNT^KO^ littermate 3T3 cell lines probed for the indicated proteins. **F**. Model of MLX activation versus repression and the role of MondoA and MNT in WT versus MLX^KO^ cells.

To further explore the potential role of MNT in apoptosis suppression in germ cells, we treated male germ cell-derived lines with siRNAs against MLX and MNT. While no effect was observed in GC-2 mouse spermatocyte-like lines treated with siRNA against MLX, MondoA or MNT (Figure S7A), siRNA against MNT in the GC-1-spg spermatogonia-like mouse germ cell line decreased viability to a similar extent as siRNAs against MLX (Figure 7A). This may be due to the elevated expression of MNT in GC-1-spg compared to GC-2 cells (Figure S7B). As observed for siRNA against MondoA or MLX previously (Figure 4E), siMNT kills NTera2 cells, derived from a human male germ cell tumor (Figure 7B). As observed in MNT^KO^ 3T3 cells, siMNT treatment of GC-1-spg cells strongly induced both TXNIP and BIM (Figure 7C). Moreover, loss of either MLX or MNT triggers cell death in the NTera2 cell line by activating similar stress pathways (BIM, p-*γ*H2AX, cleaved-PARP) as upregulated in the tissues and cells derived from MLX^KO^ animals (Figure 7E, Figure 6E, Figure S5D). Together our data support roles for both MLX and MNT in the growth and survival of multiple cell types.

**Figure 7:**
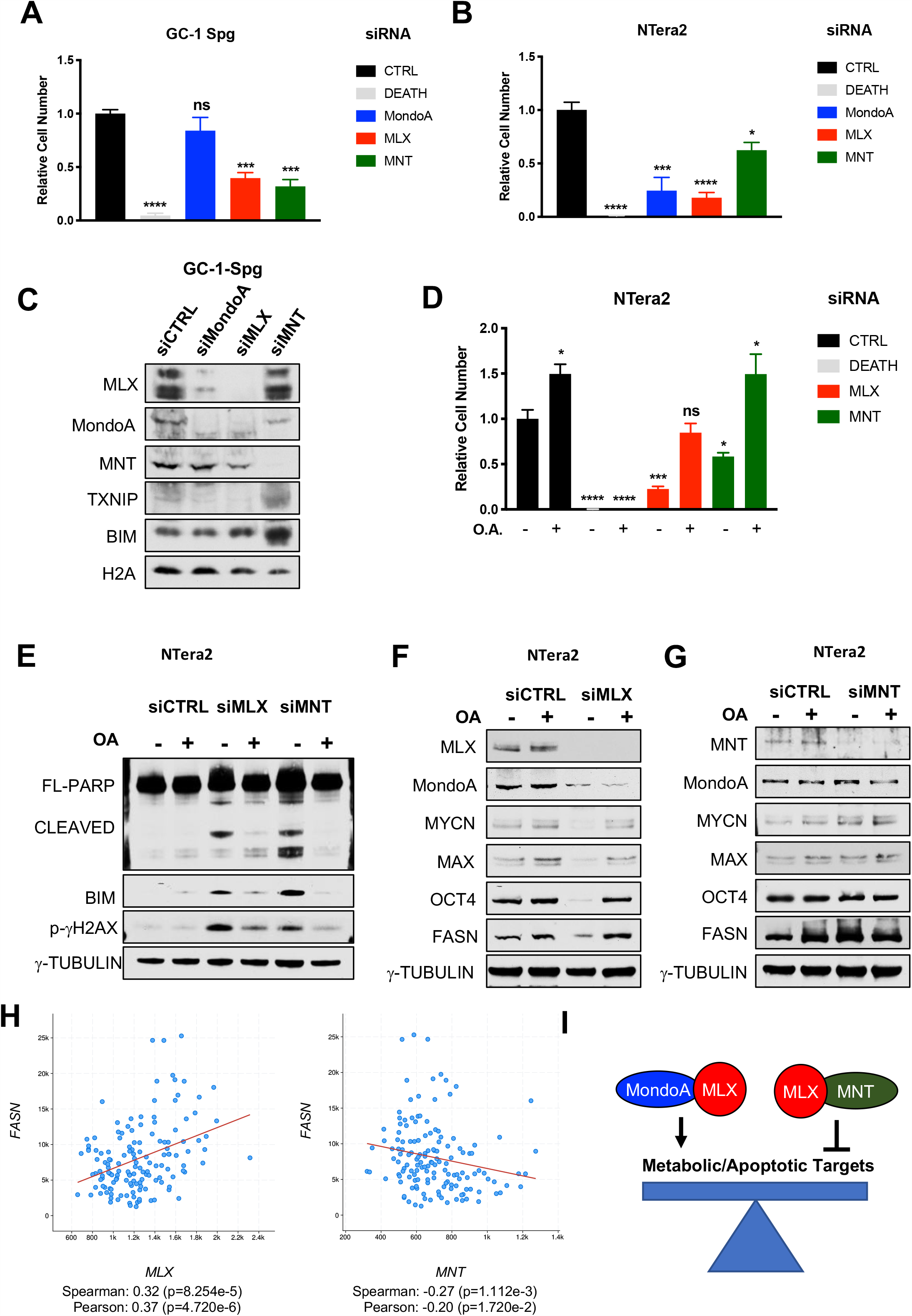
MLX and MNT regulate metabolism and survival of male germ cell lines. **A**. Relative viable cell number of the GC-1-Spg cells after siRNA transfection with the indicated siRNA with siDEATH included as a control for siRNA transfection efficacy (N=3 independent experiments, shown is the mean +/- SD). **B**. Relative viable cell number of the NTera2 cells after siRNA transfection with the indicated siRNA with siDEATH included as a control for siRNA transfection efficacy (N=4 independent experiments, shown is the mean +/- SEM). **C**. Western blot analysis of GC-1-Spg cells transfected as in (**A**) probed for the indicated proteins. **D**. Relative viable cell number of the NTera2 cells after siRNA transfection with the indicated siRNA with siDEATH included as a control for siRNA transfection efficacy. Cells were cultured in the presence or absence of 35uM oleic acid O.A. (N=4 independent experiments, shown is the mean +/- SEM). **E-G**. Western blot analysis of NTera2 cells as in (**D**) probed for the indicated proteins. **H**. Co-expression analysis of *MLX* and *FASN* and *MNT* and *FASN* from the Testicular Germ Cell Tumor Dataset (TCGA PanCancer Atlas). **I**. Model of MLX activation versus repression and the role of MondoA and MNT in these processes.

We previously reported that MondoA knockdown in MYCN-driven neuroblastoma cells leads to induction of apoptosis due, at least in part, to decreased FASN expression and attenuated fatty acid biosynthesis. The MondoA deficient neuroblastoma cells could be rescued by addition of oleic acid (a monounsaturated C18:1 fatty acid) (Carroll et al., 2015). We confirmed that this was also the case in the NTera2 cells (Figure S7D). Because MLX is a functional heterodimeric partner for MondoA we tested whether oleic acid treatment of MLX or MNT knockdown cells affected their growth and survival. As shown in Figures 7D-E, while loss of MLX decreased MYCN, MAX, OCT4 and FASN expression in NTera2 cells, oleic acid treatment reversed this effect and suppressed stress markers (Figure 7E-F). Furthermore, MNT knockdown increased FASN expression (Figure 7G and Figure S7C), while having no effect on MYCN-MAX or OCT4 levels (Figure 7G). These findings are consistent with TCGA data from male germ cell tumors where higher MLX transcript levels positively correlate with *FASN expression*, while *MNT* expression is inversely correlated with *FASN* (Figure 7H). This suggests that MNT antagonizes a subset of MondoA-MLX targets (modeled in Figure 7I).

Lastly, we examined a dataset comparing normospermic to teratozoospermic men (Platts et al., 2007) which indicated a significant correlation of male sterility with decreased levels of mRNA encoding *MLX, MNT*, as well as *CREM* and its targets (including *TNP* and *PRM* transcripts) and increased levels of *BCL2L11, FASN* and *XBP1* (Figure S7D). These findings match many of the key changes observed in the MLX^KO^ testes compared to WT (summarized in Figure S7E) These findings are consistent with a role for MNT-MLX interactions and their shared transcriptional targets in human male fertility by acting as regulators of mammalian spermatogenesis.

## Discussion

Here we describe a previously unexplored function of the MLX-based arm of the extended MYC network, namely an absolute requirement for MLX (and MondoA) in normal male germ cell (GC) development and function. Inactivation of the MLX arm of the network leads to male-specific sterility, altered metabolism, and increased stress, accompanied by activation of apoptosis in GCs, resulting in oligoasthenoteratozoospermia (OAT). This is an intriguing phenotype, as idiopathic OAT is typically associated with a variety of metabolism-related pathologies, including metabolic syndrome, diabetes, obesity and inflammation (Leisegang et al., 2019), all of which have been reported to be linked with dysregulation of MLX’s dimerization partners MondoA and/or ChREBP (reviewed in (Peterson and Ayer, 2011) (Richards et al., 2017)). While MYC, MYCN, MGA and MAX have all been shown to play distinct and critical roles in male germ cell development (summarized in Figure S7G), with MYC/MYCN (and MAX) regulating SSC function and prolifetation, whereas MAX (with MGA) represses meiosis-associated transcription (Kanatsu-Shinohara et al., 2016) (Suzuki et al., 2016), these four genes are also required for normal embryonic development (Davis et al., 1993) (Charron et al., 1992) (Stanton et al., 1992) (Shen-Li et al., 2000) (Burn et al., 2018)). By contrast, MLX and MondoA are both dispensable for embryogenesis (see Figure 1). Nonetheless, MLX deletion does lead to a loss of viability in a subset of cell types *in vivo* (male GCs) and *ex vivo* in the context of metabolic stress (e.g. splenocytes, 3T3s and tumor cells (e.g., MGCT)). This consistent with the key role of nutrient-sensing in male GC development, and suggests possible routes of dysregulation associated with idiopathic OAT in humans.

Relevant to a critical role for MLX in GC metabolism we observed that the serum and testes of MLX^KO^ mice exhibit altered metabolism consistent with a shift in glucose metabolism from oxidative phosphorylation, via the TCA cycle, to the production of lactate, and to oxidation of alternative TCA substrates, such as branched-chain amino acids, and fatty acids. Deletion of MondoA in mice, leads to a similar change in serum metabolites, associated with enhanced glycolysis and activation of beta-oxidation (Imamura et al., 2014) (Ahn et al., 2016)). Moreover, loss of the MondoA-MLX target TXNIP is sufficient to induce a similar metabolic profile, including augmented CPT1A expression and activity (Chen et al., 2016). This suggests that the choice of oxidative substrate for the TCA cycle in testes is controlled by MondoA-MLX through its regulation of TXNIP. These metabolic alterations, observed in the MLX^KO^ testes, are associated with GC apoptosis, concomitant with aseptic inflammation and immune cell activation, symptoms also associated with OAT.

A recent study in humans linked a SNP in *MLX* to hyper-activation of TXNIP in Takayasu Arteritis (Tamura et al., 2018), which is also an inflammatory disorder. Interestingly, TXNIP is both a mediator of inflammasome activation and ER stress (Shalev, 2008) (Chen et al., 2010)) but MEFs derived from TXNIP^KO^ embryos exhibit a constitutively activated ER stress state, rather than an attenuated one (similar to what we observe in both MLX^KO^ or MNT^KO^ 3T3 cells, Figure 6E) (Lee et al., 2014). However, in contrast to our findings with MLX or MondoA deletion, loss of TXNIP does not result in male sterility (Oka et al., 2006). This suggests that critical MLX transcriptional targets other than TXNIP are responsible for the majority of the male-specific sterility phenotype associated with loss of either MLX or MondoA.

ChREBP is another dimerization partner of MLX, and its loss of function has been previously shown to result in decreased multi-tissue glycolysis and lipogenesis (Iizuka et al., 2004), features which are not recapitulated in our MLX null mice. This may be indicative of a predominant role for MondoA-MLX compared to ChREBP-MLX in whole-body metabolism, as expected given the more tissue-restricted expression of ChREBP (e.g. liver). Nonetheless, altered myeloid cell function and impaired immune response is observed in ChREBP^KO^ mice (Sarrazy et al., 2015)), and would also be expected to be MLX-dependent.

Our transcriptional profiling and genomic occupancy analyses using ChIP-Seq has identified direct targets (i.e. genes bound and regulated) of MLX in the testes, including metabolic and stress effectors as well as many genes relevant to male-specific germ cell development. Among the latter are 6% of the approximately 5800 genes previously known to be bound by the essential transcriptional regulator of spermiogenesis, CREM. These include protamines and transition proteins. These data strongly support the notion that MLX and its binding partners act as transcriptional mediators of critical events in mammalian GC development in concert with other transcriptional regulators such as CREM and MAX. Interestingly, neither MLX nor MAX occupy a significant number of loci that are targets of the DMRT1 testes-specific transcriptional regulator of differentiation. Thus, the direct activity of MLX and other MYC network members in GC development may, to some extent, be more focused on functions regulated through the CREM pathway. Among shared targets are gene products that are responsible for the metabolic, stress-related, and apoptotic phenotypes observed in MLX^KO^ testes such as increased abundance of acylated carnitine, CPT1A, FAS, and BIM.

Many of the MLX-linked metabolic and stress targets observed in testes are not cell-type specific and we find their expression altered in both primary lymphocytes and in 3T3 cells derived from WT vs MLX^KO^ embryos. MLX loss of function sensitizes these cells to apoptotic stimuli and effectors known to be dependent upon MYC (e.g. FASL-FAS interactions, BIM) (Hueber et al., 1997) (Muthalagu et al., 2014). Taken together, this data supports a direct role for MLX in both regulating important metabolic targets (as are known to be regulated by MondoA-MLX and ChREBP-MLX) but also direct transcriptional repression of apoptosis effectors (such as BIM), most likely mediated by MLX heterodimerization with repressors such as MNT. Indeed, our occupancy analyses indicate that genes bound by MLX are also bound by other members of the MYC network (such as MAX and MNT). Loss of MLX appears to shift the occupancies of these other factors: for example, MAX and MNT bind to a large number of previously unoccupied sites. Such shifts in binding upon perturbation of the network could potentially lead to dysregulation of the extended network triggering, in turn, a stress and apoptotic response. We hypothesize that formation of transcriptional condensates, as recently described for other transcriptional networks (reviewed in (Sabari et al., 2020)), may underlie the apparent multiprotein occupancy we observe for MLX and other MYC network proteins.

Our study further implicates the extended MYC network, and specifically the nutrient-sensing MLX arm, in the direct regulation of, and linkage between, differentiation, metabolism and apoptosis. Importantly, metabolic programs change along with changes in cellular state, and thus must be responsive to both extrinsic signals, such as mitogenic cues (via effectors such as MYC-MAX), but also to intrinsic metabolic cues (as in the case of G6P and acidosis, known to activate MondoA-MLX (Wilde et al., 2019)), or lipid droplets known to inhibit MondoA-MLX activity) (Mejhert et al., 2020). It is clear that functional interactions among MYC network members are functionally relevant, not only in MYC-driven oncogenesis, but also during MGC development (this work), and regeneration of both skeletal muscle and liver (Hunt et al., 2015) (Wang et al., 2018)). We anticipate that further genetic perturbation of the network in distinct biological contexts, coupled with high-resolution genomic analysis will yield important insights into the molecular control of normal and abnormal cellular behavior in both tissue homeostasis and oncogenesis.

## Materials and Methods

### Animal Use

All experiments involving mice were carried out under accordance with the guidelines of the following institutions: Ethics Statement: This study was performed in strict accordance with the recommendations in the Guide for the Care and Use of Laboratory Animals of the National Institutes of Health. All of the animals were handled according to approved institutional animal care and use committee (IACUC) protocol 50783 of the Fred Hutchinson Cancer Research Center. The animal experiments conducted at the University of Utah were performed under IACUC protocol 15-04012. Every effort was made to minimize pain and suffering.

### Generation of *Mlx*^*-/-*^ mice

The *Mlx* knockout allele was created in our lab by a homologous recombination targeted method in 129S4 AK7 murine ESCs. The targeting construct is in the PGKneoF2L2DTA backbone and is based upon the coding sequence of *Mlx* transcript variant 1 (encoding MLX-*α* protein). This construct includes in 5’-to-3’ order: **1)** a 1546 bp 5’-homology arm including exons 1-2 of *Mlx*, **2)** a 1717 bp Loxp-flanked region encoding exons 3-6 (bHLHLZ domain) of *Mlx*, **3)** a FRT-flanked PGKNEO-positive selection cassette, **4)** a 2873 bp 3’ homology arm spanning exon 7 of *Mlx* transcript variant 1 and a pGKDTA-negative selection cassette. Selected ESC clones were injected into blastocysts to generate chimeric animals. These chimeras were bred to ROSA26 FlpO/FlpO females (Raymond and Soriano, 2007) in the 129S4 co-isogenic background to remove the Frt-flanked NEO cassette to get conditional knockout mice or Meo2-Cre delete mice (Tallquist and Soriano, 2000) to get total knockout mice. The FlpO allele or the Meox2-Cre allele was subsequently crossed out. Two independent mouse lines from independent ESC clones were found to be phenotypically indistinguishable. All the targeting mice were confirmed by PCR and Southern blotting with 5’ external, 3’ external and NEO probes.

### Genotyping

The following primers were used as a mixture of 3 primers for PCR to discriminate WT from KO from floxed animals; MLX forward (MLX 5644f): 5’ actccaggaaaagtgtagctgcc 3’, MLX reverse (MLX 5845r): 5’ caagctgttggcttccatacagg 3’, MLX deletion (MLX 3679f): 5’ caaccatggtcacacctggttc 3’ yielding the following size PCR products: MLX forward + MLX reverse: WT=201bp, flox=327bp MLX deletion + MLX reverse: KO=589bp. The annealing temperature used was 65°C.

### Mating Tests

Pairs of sexually mature mice (one of each sex) were housed in the same cage and mating was observed with the seminal plugs and number of pups born recorded. For Mendelian frequency, heterozygotes were bred with heterozygotes and the progeny (F1) were genotyped. For knockout (KO) mating tests, homozygous null males or females were bred with WT mice.

### Metabolomics

### Reagents

Acetonitrile, ammonium acetate, and acetic acid (LC-MS grade) were all purchased from Fisher Scientific (Pittsburgh, PA). The standard compounds corresponding to the measured metabolites were purchased from Sigma-Aldrich (Saint Louis, MO) and Fisher Scientific (Pittsburgh, PA).

### Serum sample preparation and LC-MS/MS measurement

Male mice (3 of each genotype WT, MLX^KO^) were bled by retroorbital eye-bleed and serum was isolated, then flash-frozen on dry ice before subsequent use for LC-MS/MS. Frozen serum samples were first thawed overnight at 4°C, and 50 μL of each sample was placed in a 2 mL Eppendorf vial. The initial step for protein precipitation and metabolite extraction was performed by adding 500 μL MeOH and 50 μL internal standard solution (containing 1,810.5 μM ^13^C_3_-lactate and 142 μM ^13^C_5_-glutamic acid). The mixture was then vortexed for 10 s and stored at −20 °C for 30 min, followed by centrifugation at 14,000 RPM for 10 min at 4 °C. The supernatants (450 μL) were was collected into a new Eppendorf vial, and dried using a CentriVap Concentrator (Labconco, Fort Scott, KS). The dried samples were reconstituted in 150 μL of 40% PBS/60% ACN. A pooled sample, which was a mixture of all serum samples was used as the quality-control (QC) sample. The LC-MS/MS experimental procedures were well documented in our previous studies (Zhu et al., 2014) (Barton et al., 2015) (Carroll et al., 2015) (Gu et al., 2015) (Gu et al., 2016; He et al., 2020; Shi et al., 2019)). Briefly, all LC-MS/MS experiments were performed on a Waters Aquity I-Class UPLC-XenoTQ-S micro (Waters, MA) system. Each sample was injected twice, 10 µL for analysis using negative ionization mode and 2 µL for analysis using positive ionization mode. Both chromatographic separations were performed in hydrophilic interaction chromatography (HILIC) mode on a Waters XBridge BEH Amide column (150 x 2.1 mm, 2.5 µm particle size, Waters Corporation, Milford, MA). The flow rate was 0.300 mL/min, auto-sampler temperature was kept at 4 °C, and the column compartment was set at 40 °C. The mobile phase was composed of Solvents A (5 mM ammonium acetate in 90%H_2_O/ 10% acetonitrile + 0.2% acetic acid) and B (5 mM ammonium acetate in 90%acetonitrile/ 10% H_2_O + 0.2% acetic acid). After the initial 2 min isocratic elution of 90% B, the percentage of Solvent B was linearly decreased to 50% at t=5 min. The composition of Solvent B maintained at 50% for 4 min (t=9 min), and then the percentage of B was gradually raised back to 90%, to prepare for the next injection. The mass spectrometer is equipped with an electrospray ionization (ESI) source. Targeted data acquisition was performed in multiple-reaction-monitoring (MRM) mode. We monitored 121 and 80 MRM transitions in negative and positive mode, respectively (201 transitions in total). The whole LC-MS system was controlled by MassLynx software (Waters, MA). The extracted MRM peaks were integrated using TargetLynx software (Waters, MA).

### Cell sample preparation and LC-MS/MS measurement

Isolated cells from the seminiferous tubules of age-matched WT and MLX^KO^ mice (N=3) were separated into 4 technical replicates of 1×10^6 cells each and flash frozen on dry ice. Soluble metabolites were extracted into 1 ml of 20:80% water:methanol before clearing the insoluble and drying down on a Speed-Vac before subsequent LC-MS/MS. The LC-MS/MS experiments were performed on an Agilent 1260 LC-6410 QQQ-MS (Agilent Technologies, Santa Clara, CA) system. 5 µL of each sample was injected for analysis using positive ionization mode. Chromatographic separation was performed using a Waters XSelect HSS T3 column (2.5 µm, 2.1 x 150 mm). The flow rate was 0.3 mL/min. The mobile phase was composed of Solvents A (100% H_2_O with 0.2% formic acid) and B (100% ACN with 0.2% formic acid). After the initial 0.5 min isocratic elution of 100% A, the percentage of Solvent A was linearly decreased to 5% at t=10 min. Then the percentage of A remained the same (5%) for 5 min (t=15 min). The metabolite identities were confirmed by spiking with mixtures of standard compounds. The extracted MRM peaks were integrated using Agilent Masshunter Workstation software (Agilent Technologies, Santa Clara, CA).

### Serum Testosterone Quantification

Blood was collected by either retro-orbital eye bleed, or cardiac stick and heparinized plasma was isolated. Testosterone levels were determined by radioimmunoassay at the University of Virginia Center for Research in Reproduction (NICHD Grant # U54-HD028934) Charlottesville VA.

### Cell Isolation from Testicular Tissue

Testes and epididymides were dissected, defatted, and processed as follows for cellular isolation: for seminiferous tubule cell isolation, testes were dissected, decapsulated, and subjected to an enzymatic digestion to isolate seminiferous tubules from the interstitial stromal cells (including Ledig and immune cells). Isolated tubules were then digested to release a single cell suspension of the seminiferous epithelium including germ cells and Sertoli cells. For epididymal germ cells, epididymides were dissected and the caudal portion was cut to release cellular content of sperm. Single cell suspension was filtered to remove tissue yielding predominately mature spermatozoa from WT tissue.

### Spermatogenesis Analysis

Testes and epididymides were dissected, defatted, and processed for analysis as previously described (Amory et al., 2011). For calculation of testicular daily sperm production rate (spermatid count), testes were weighed and homogenized in 0.1M sodium phosphate buffer (pH 7.4) with 0.1% Triton X-100 via eight strokes of a 15 ml Kontes homogenizer. Homogenization-resistant spermatids were counted on a hemocytometer and spermatid per gram per testis were calculated (Amann, 1986).

### Cell Culture

3T3 cell lines, Ntera2, HepG2, GC-1-Spg and GC-2-Spd(ts) cells were all cultured in DMEM with 10% FBS and penn/strep. Primary B220+ sorted splenic B cells were cultured in RPMI with 15% FBS, penn/strep, beta-mercaptol, and LPS (1ug/ml from Sigma-Aldrich).

### RNAi Transfection

Flexitube Gene Solutions siRNA mixtures (Qiagen) were utilized to knockdown the indicated target. RNAiMax (Qiagen) was used according to the manufacture’s conditions. Cells were counted 72-96 hours post-transfection and viability was monitored by trypan blue exclusion. For oleic acid treatment, Oleic Acid Water Soluble (Sigma-Aldrich) was resuspended in sterile water.

### B220+ Cell Purification

Spleens were smashed and B220+ B cells were purified from the splenocytes using AutoMACS system according to the manufacturers recommendations. Purity was routinely over 95% pure, as assessed by Flow Cytometry.

### Flow Cytometry

Testis tissue was prepared as in (Bastos et al., 2005). Isolated seminiferous tubule or epididymal cells were resuspended in staining buffer with HOECHST and samples were run on a Canto-2 Flow Cytometer and analyzed by FACS-Diva Software.

### Immunohistochemistry Tissue Staining

Testicular and epididymal tissue was fixed in Modified Davidson’s Fluid (MDF) as described(Latendresse et al., 2002), then embedded in paraffin and sectioned onto slides at 5 micron thickness. Slides were deparaffiinized, rehydrated and antigen-retrieval was utilized. For Immunofluorescent IHC, Dako reagents were used: (Block, Primary Dilution Buffer, Antifade mounting media), and Alexa-Flour 488nm secondary antibodies were used in combination with DAPI for staining. For tissue staining and IHC, samples prepared as described above were submitted to the FHCRC Experimental Histopathology Core and either stained with hematoxylin and eison, or stained with the indicated antibodies, visualized with the cromophore DAB and counterstained with hematoxylin to mark nuclei.

The staining of previously published Xenograft tissue of the NTera2 cells (Venkataramani et al., 2012) were carried out by the Institute of Pathology of the University Medical Center Göttingen. Briefly, 4 micron thick sections were mounted on slides, deparaffinnized, rehydrated, and antigen-retrieval was utilized. The slides were stained with primary antibody then biotinylated secondary antibodies using a REAL Detection System (LSAB+ kit; Dako). The signals were visualized using a REAL Streptavidin Alkaline Phosphatase kit (Dako), while Ki-67 staining was visualized with DAB. All samples were counterstained with hematoxylin, mounted in super mount medium, and analyzed via light microscopy.

### Western Blots

Tissues and cell pellets were lysed in RIPA buffer. For whole testes sample preps, tissue was homogenized mechanically to facilitate lysis. Lysates were quantified by BCA assay (Pierce) or normalized to cell number for equal loading. Samples were resolved on NuPAGE 4-12% Bis-Tris gradient gel before transferring to Nitrocellulose (0.2 micron). Blots were blocked with 5% Milk in TBST, washed with TBST, probed with primary and secondary antibody in 5% Milk in TBST. The secondary antibody was HRP conjugated and chemiluminescent detection was employed. Blots were exposed to Pro-Signal Blotting Film (Genesee Scientific).

### RNA Extraction and Sequencing

RNA was extracted with Trizol reagent, quantified on a TapeStation. 500ng of RNA was submitted for library preparation through FHCRC Genomics Core. Libraries were aligned to mm10 using TopHat then processed with EdgeR or DE-Seq. Data was analyzed with GSEA and volcano and violin plots were generated using ggplot.

### Chromatin-Immunoprecipitation and Sequencing (ChIP-Seq)

We performed chromatin immunoprecipitation and sequencing (ChIP-seq) as previously described (Skene and Henikoff, 2015) with modifications to improve solubility of transcription factors, which tended to vary depending on cell type. The chromatin preparations from the testes were from a pool of testes from 6 WT/KO animals, and this material was not treated with MNase. The chromatin preparations from the B cells and 3T3 cell lines were treated with MNase. Briefly, after formaldehyde cross-linking, cell lysis, and chromatin fragmentation with MNase, the final SDS concentration after dilution of total chromatin was increased to 0.25% with addition of 20% SDS stock solution. Sonication was performed in a Covaris M220 focused ultrasonicator for 12 minutes with the following settings: 10% duty cycle, 75W peak incident power, 200 cycles/burst, and 6-7°C bath temperature. The SDS concentration of the sonicated chromatin solution was readjusted to 0.1% with dilution buffer. Immunoprecipitation was performed on the clarified chromatin (input) fraction from 10×10^6 cellular equivalents and DNA was purified was using standard phenol:chloroform extraction. The following antibodies were used for ChIP: MAX C-17/sc-197 (Santa Cruz Biotechnology, Cat. No. sc-8011X), MLX D8G6W (Cell Signaling Technology, 85570S), MNT (Bethyl, A303-627A), and the negative control GFP (Cell Signaling Technology, 2956S). We used 10 μg of antibody for each immunoprecipitation. To purified ChIP DNA, we added 10 pg of spike-in DNA purified from MNase-digested chromatin from *D. melanogaster* S2 cells or *S. cerevisiae* (Skene and Henikoff, 2017) to permit comparison between samples. DNA was then subjected to library preparation as previously described (Henikoff et al., 2011) (Orsi et al., 2015)and 25×25 paired-end sequencing was performed on an Illumina HiSeq 2500 instrument at the Fred Hutchinson Cancer Research Center Genomics Shared Resource. For the B cells samples, an alternative library preparation was employed (Ramani et al., 2019).

Sequencing datasets were aligned to the mouse mm10 genome assembly using Bowtie2. Datasets were also aligned to the dmel_r5_51 (*D. melanogaster*) or sc3 *(S. cerevisiae*) assemblies using Bowtie2 depending on the source of the spike-in DNA. Counts per base-pair were normalized as previously described by multiplying the fraction of mapped reads spanning each position in the genome by genome size (Kasinathan et al., 2014) or by scaling to spike-in DNA (Skene and Henikoff, 2017). Peaks were called using MACS. Plots were generated with ngs plot (Shen et al., 2014), or the R package ggplot2 Motif enrichment was done using HOMER (Heinz et al., 2010).

### Antibodies Used

The following antibodies were used for the indicated techniques with catalog number and supplier listed: anti-DDX4 (WB and IF, AB13840, Abcam), anti-Phospho-H3(ser10) (IF, 9706S, Cell Signaling Technology), anti-Phospho-gamma-H2AX (WB and IF, AB11174, Abcam), anti-Histone H2A (WB, 2578S, Cell Signaling Technology), anti-TXNIP (WB and IF, K0204-3, K0205-3, MBL International), anti-PGK1/2 (WB and IF, SC-28784, Santa Cruz Biotechnology), anti-Ki-67 (IHC, AB16667, Abcam), anti-FAS (WB and IHC, SC-1024, Santa Cruz Biotechnology), anti-TIMP1 (IHC, SC-6832, Santa Cruz Biotechnology), anti-MLX (WB, IHC and ChIP, 85570, Cell Signaling Technology), anti-MondoA (WB and IHC, 13614-1-AP, Proteintech), anti-ChREBP (WB, NB400-135, Novus), anti-MYC (WB, 13987, Cell Signaling Technology), anti-MYC (IHC, SC-764, Santa Cruz Biotechnology), anti-MYCN (WB and IHC, SC-53993, Santa Cruz Biotechnology), anti-MAX (WB, IHC and ChIP, SC-197, Santa Cruz Biotechnology), anti-MNT (WB and ChIP, A303-627A, Bethyl), anti-EOMES (WB, AB23345, Abcam), anti-OCT4 (WB and IHC, AB184665, Abcam), anti-CPT1A (WB, AB176320, Abcam), anti-IGFBP3 (WB, 10189-2-AP, Proteintech), anti-BIM (WB, 2933S, Cell Signaling Technology), anti-PARP (WB, 9542T, Cell Signaling Technology), anti-SCD (WB, AB19862, Abcam), anti-FASN (WB, AB128856, Abcam), anti-TOMM20 (WB, AB56783, Abcam), anti-ATF4 (WB, 11815S, Cell Signaling Technology), anti-Gamma-Tubulin (WB, T5326, Sigma-Aldrich), anti-Beta-Actin (WB, A5441, Sigma-Aldrich), anti-Mouse HRP (WB, 7076, Cell Signaling Technology) and anti-Rabbit HRP (WB, 7074, Cell Signaling Technology).

### Statistical Analysis

Graphpad Prism was used for all statistical data analysis unrelated to NGS datasets. P-values were calculated by Student’s T-test or One-Way ANOVA, when appropriate. For metabolomics analysis, MetaboAnalyst (Xia et al., 2009) was used. For RNA-Seq and ChIP-Seq analysis R Studio and Python were used.

## Acknowledgments

The authors are grateful to former and current members of the Eisenman Lab who have contributed to the discussion and direction of this project, as well as to the greater community that comprises the collaborative environment at the Fred Hutch and University of Washington. We also thank the Experimental Histopathology and Genomics shared resources at Fred Hutch.

## Author contributions

Patrick Carroll: Conceptualization; Investigation; Writing - original draft; Writing - review and editing

Pei Feng Cheng: Investigation; Methodology

Brian Freie: Data curation; Formal analysis

Sivakanthan Kasinathan: methodology, formal analysis

Haiwei Gu: conceptualization, investigation, formal analysis

Theresa Hedrich: Investigation

James A Dowdle: conceptualization

Vivek Venkataramani: conceptualization, investigation, methodology

Vijay Ramani: conceptualization

Daniel Raftery: Supervision, funding acquisition

Jay Shendure: supervision

Donald E. Ayer: supervision, funding acquisition

Charles H. Muller: supervision, conceptualization, investigation, methodology

Robert N. Eisenman: Project administration; Conceptualization; Supervision; Funding acquisition; Writing - original draft; Writing - review and editing

---

**Supplemental Figure 1.**
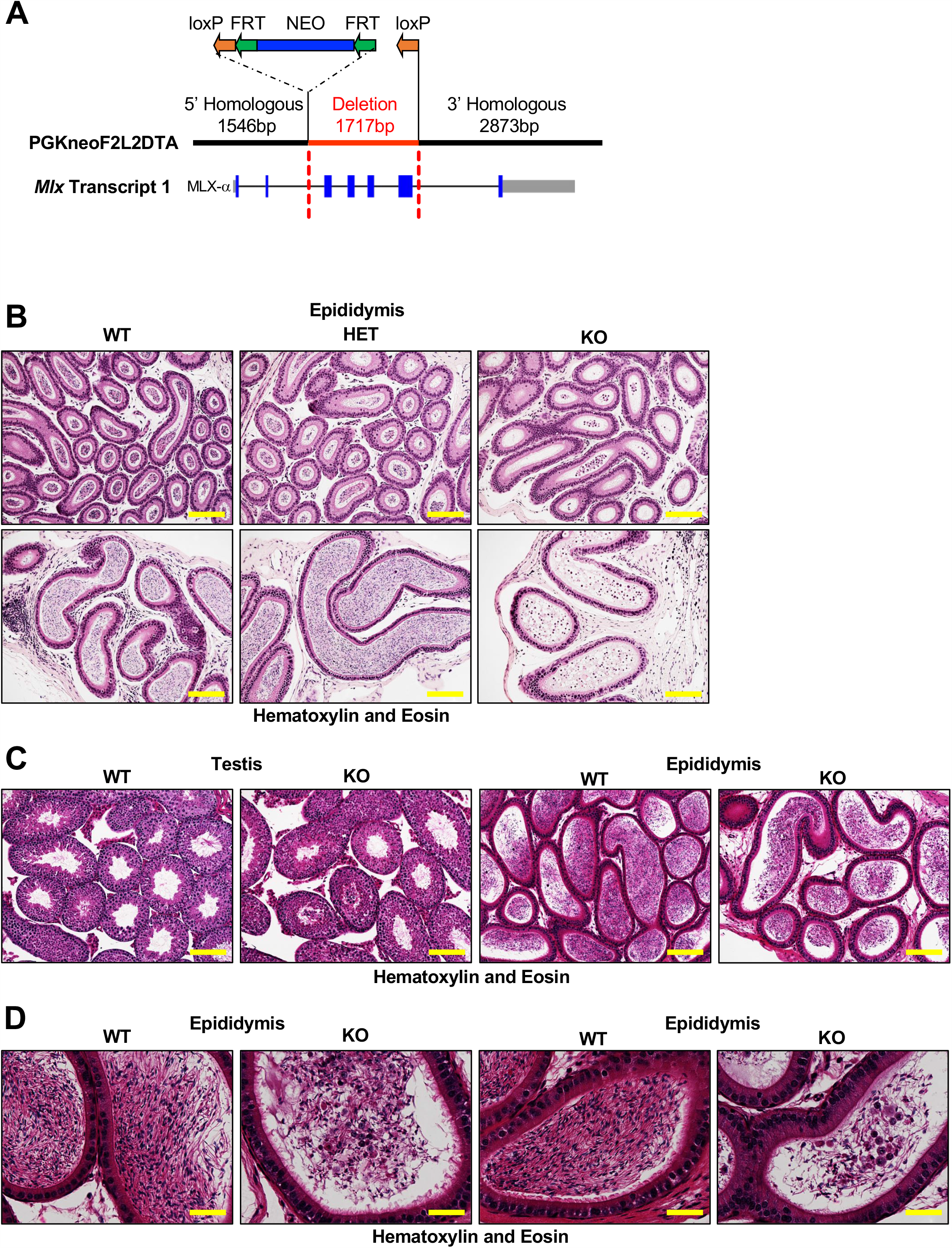
(goes with Figure 1) Histological characterization of the WT and HET versus MLX^KO^ testes and epididymides. **A**. Schematic of the targeting construct used in the deletion of murine *Mlx*. **B**. Histological analysis of WT, HET and MLX^KO^ epididymis stained with hematoxylin and eosin (100x, scale bar = 400uM). **C**. Histological analysis of P51 WT versus MLX^KO^ testis and epididymis stained with hematoxylin and eosin (100x, scale bar = 400uM). **D**. Histological analysis of P51 WT versus MLX^KO^ epididymis stained with hematoxylin and eosin (400x, scale bar = 100uM).

**Supplemental Figure 2.**
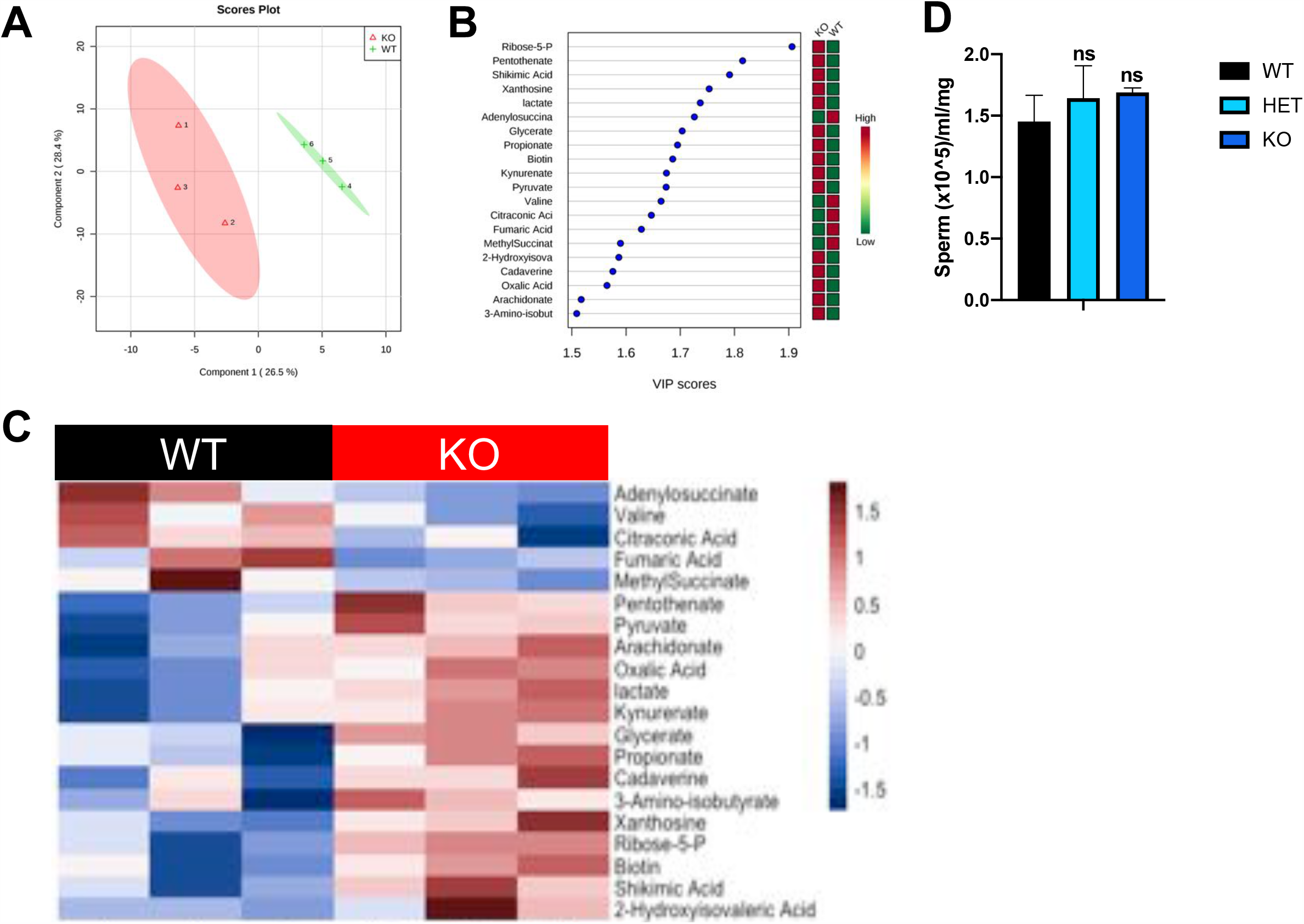
(goes with Figure 2) Metabolomic data from WT versus MLX^KO^ serum. **A**. Partial Least Square Discriminant Analysis (PLS-DA) and **B**. Variable Importance of Projection (VIP) plot from the metabolomic dataset analyzed by Metaboanalyst 4.0 (Xia et al., 2009)). **C**. Heatmap of mean-centered serum metabolomics dataset from WT versus MLX^KO^ mice showing the top 20 VIP features from PLS-DA (N=3). **D**. Sperm count from MondoA WT versus KO mice (N=5).

**Supplemental Figure 3.**
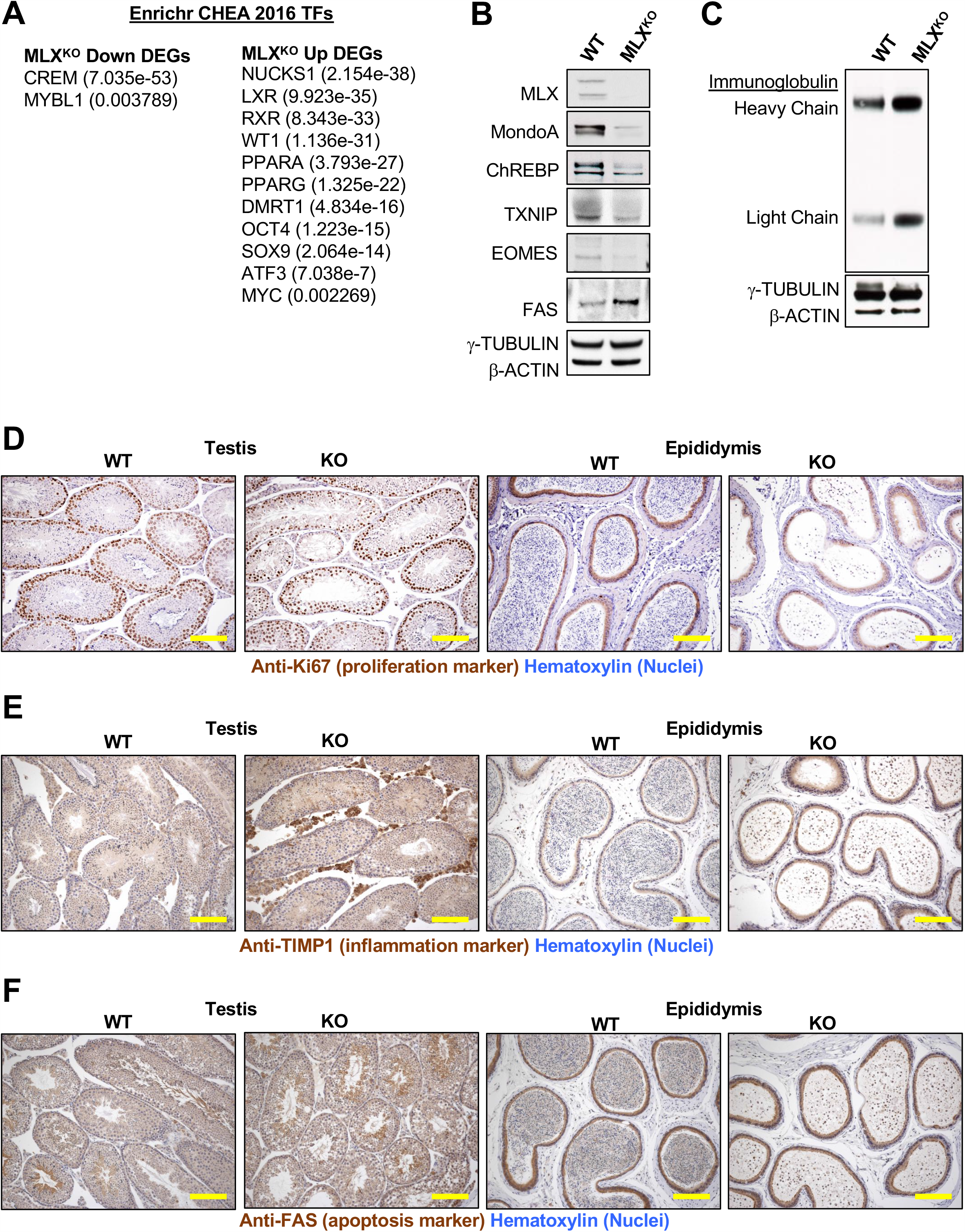
(goes with Figure 3) RNA and protein profiling of testes from WT versus MLX^KO^ mice. **A**. ChIP Set Enrichment Analysis (CHEA 2016 from Enrichr database) adjusted p-values for indicated transcription factor targets associated with Up or Down in the WT versus the MLX^KO^ RNA-Seq data. **B-C**. Western blot data from whole testes lysates from WT versus MLX^KO^ mice probed for the indicated proteins. **D**. IHC analysis of WT versus MLX^KO^ testis and epididymis stained for the proliferation marker Ki-67 (100x, scale bar = 400uM). **E**. IHC analysis of WT versus MLX^KO^ testis and epididymis stained for TIMP1 (100x, scale bar = 400uM). **F**. IHC analysis of WT versus MLX^KO^ testis and epididymis stained for FAS (100x, scale bar = 400uM).

**Supplemental Figure 4.**
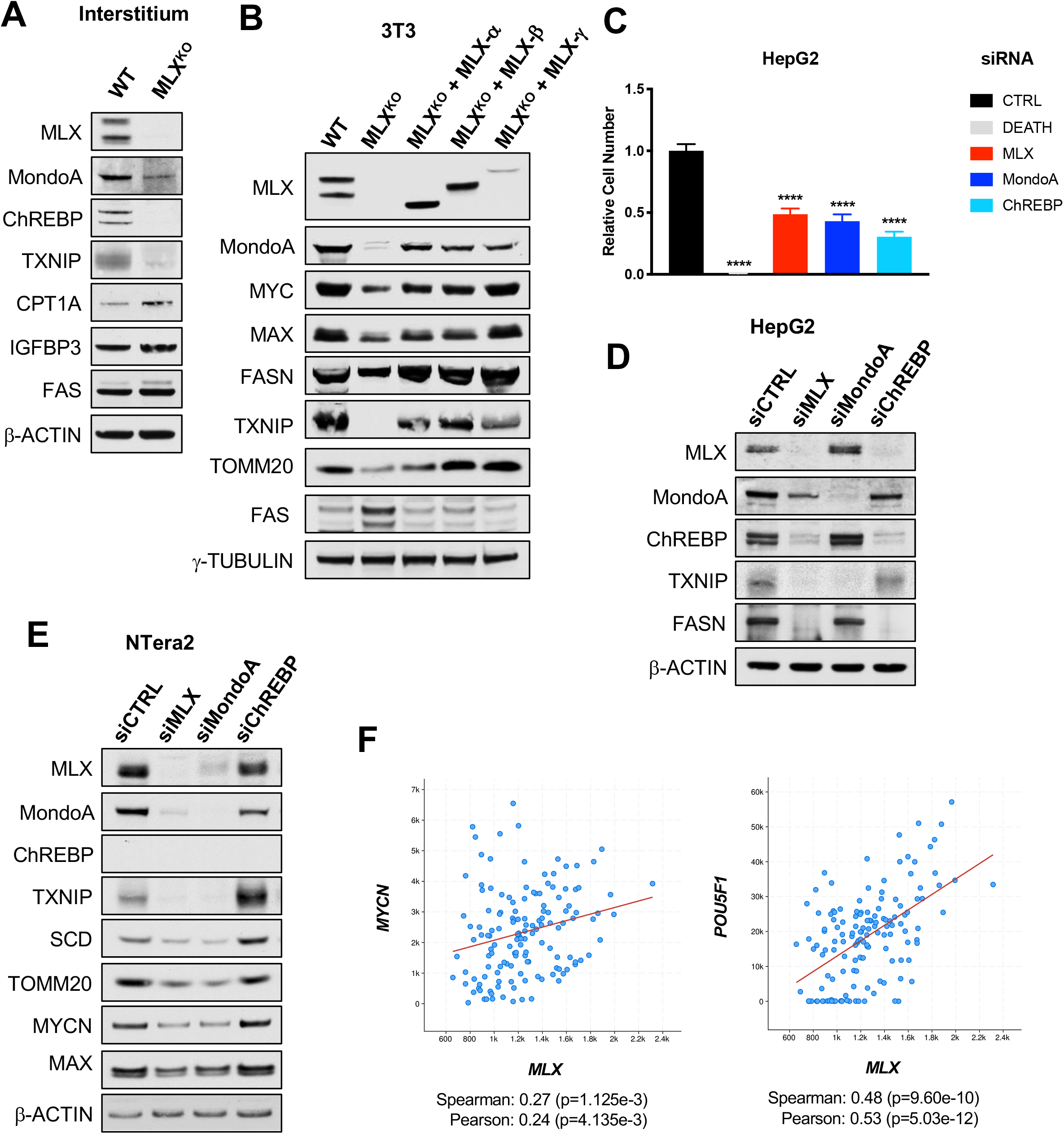
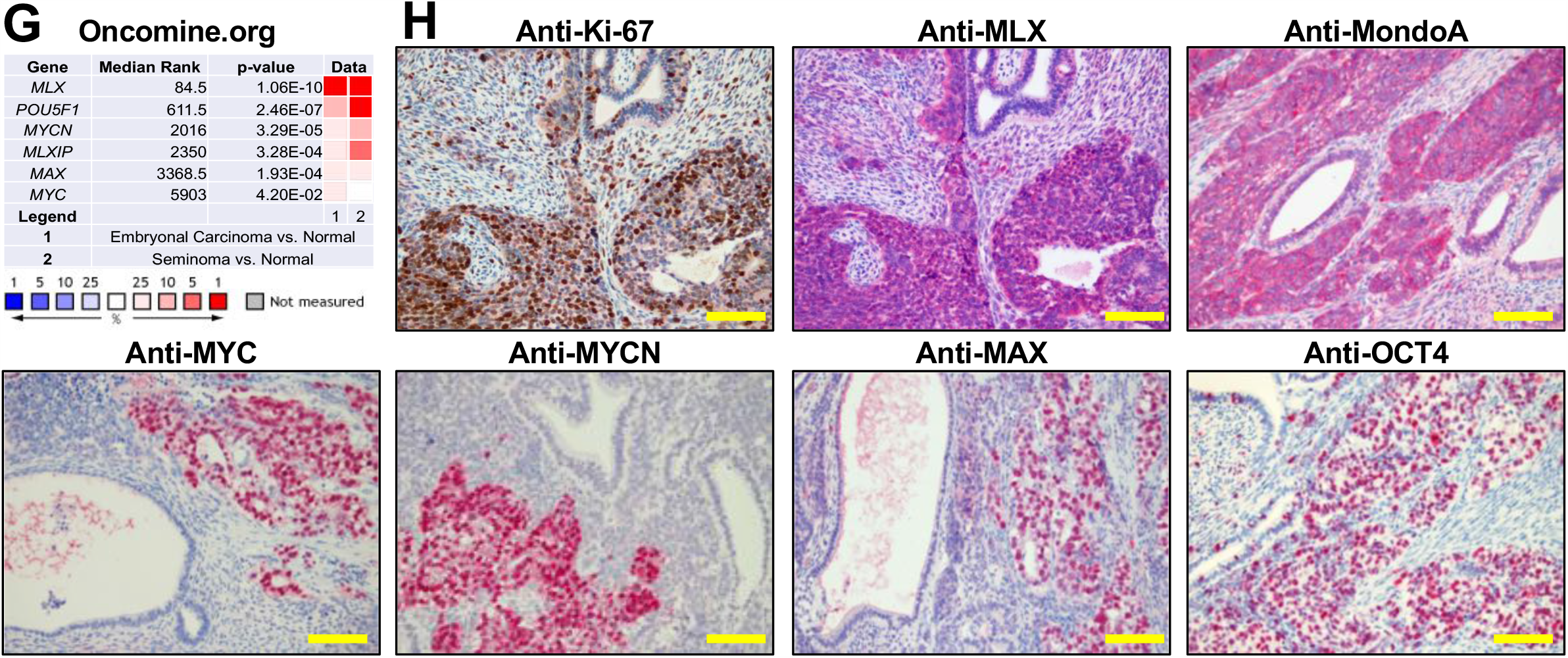
(Goes with Figure 4) Molecular, biochemical and functional validation of GSEA categories from WT versus MLX^KO^ mice. **A**. Western blot of cells isolated from the interstitium of WT versus MLX^KO^ testes tissue probed for the indicated proteins. **B**. Western blot analysis of the WT versus MLX^KO^ 3T3 cell lines reconstituted with empty vector or the indicated isoform of MLX probed for the indicated proteins. **C**. Relative viable cell number of HepG2 cells after siRNA transfection with the indicated siRNA with siDEATH included as a control for siRNA transfection efficacy (N=4 independent experiments, shown is the mean +/- SEM). **D**. Representative western blots of the HepG2 cells treated with the indicated siRNA as in (**C**.). **E**. Western blot analysis of the NTera2 cells treated with the indicated siRNA, probed for the indicated proteins. **F**. Co-expression analysis of *MLX* and *POU5F1* from the Testicular Germ Cell Tumor Dataset (TCGA PanCancer Atlas). **G**. Overexpression of the indicated mRNAs from Oncomine.org Korkola et al dataset. **H**. IHC analysis of NTera2 cell xenograft from (Venkataramani et al., 2012) stained for the indicated proteins (200x, scale bar = 200uM).

**Supplemental Figure 5:**
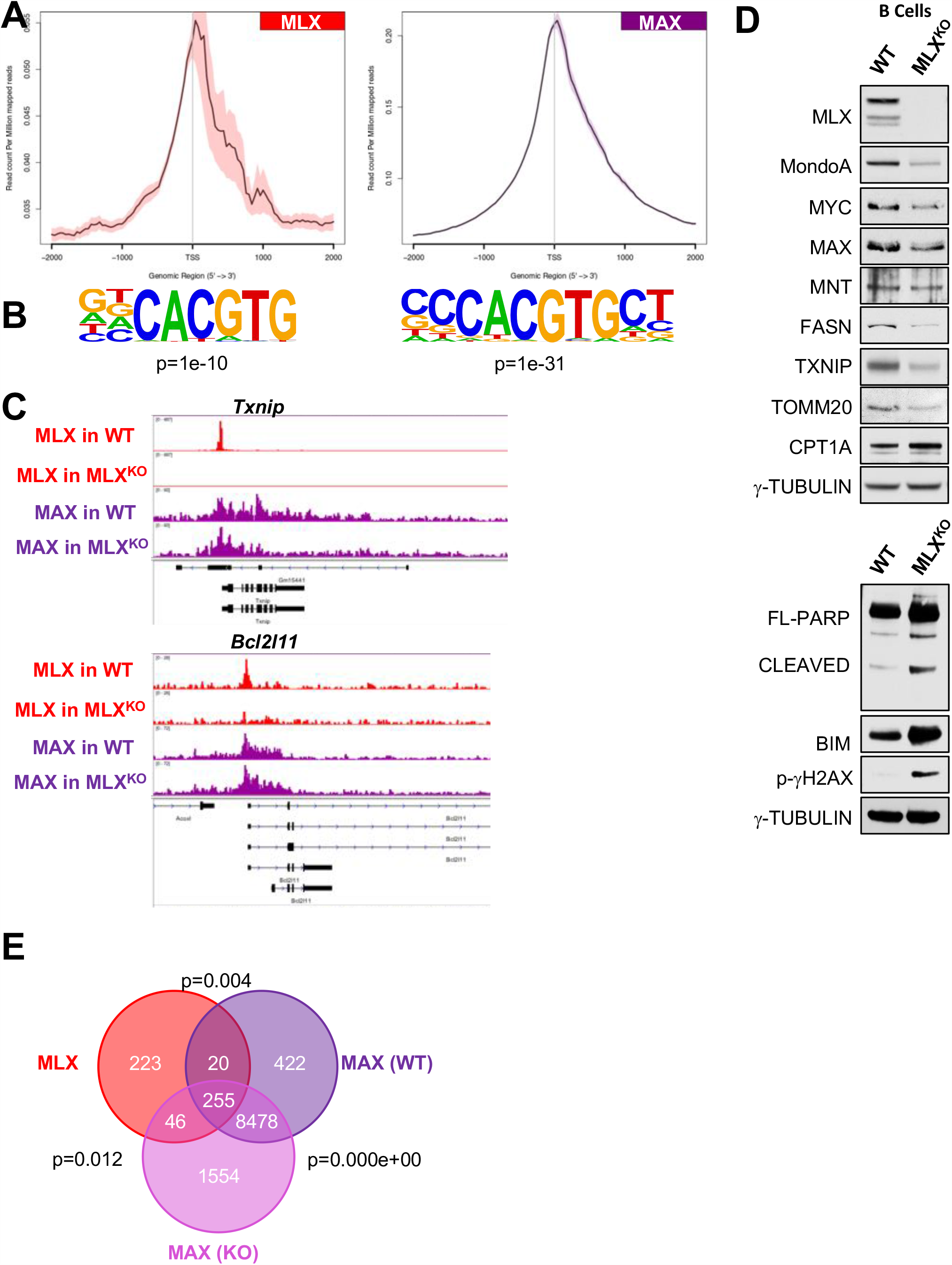
(Goes with Figure 5) MLX genomics in primary B cells shares features with the testes. **A**. NGS plots for MLX and MAX generated from ChIP-Seq data from B220+ splenic B cells cultured for 72 hours in LPS. **B**. E-Box motif from HOMER for MLX and MAX. **C**. IGV tracks from WT and MLX^KO^ B cell ChIP-Seq data for *Txnip* and *Bcl2l11 (Bim)*. **D**. Western blot analysis of the indicated proteins from WT and MLX^KO^ B cells cultured for 72 hours in LPS. **E**. Venn diagram of genes bound by MLX and MAX in WT and MLX^KO^ B cells identified by ChIP-Seq.

**Supplemental Figure 6:**
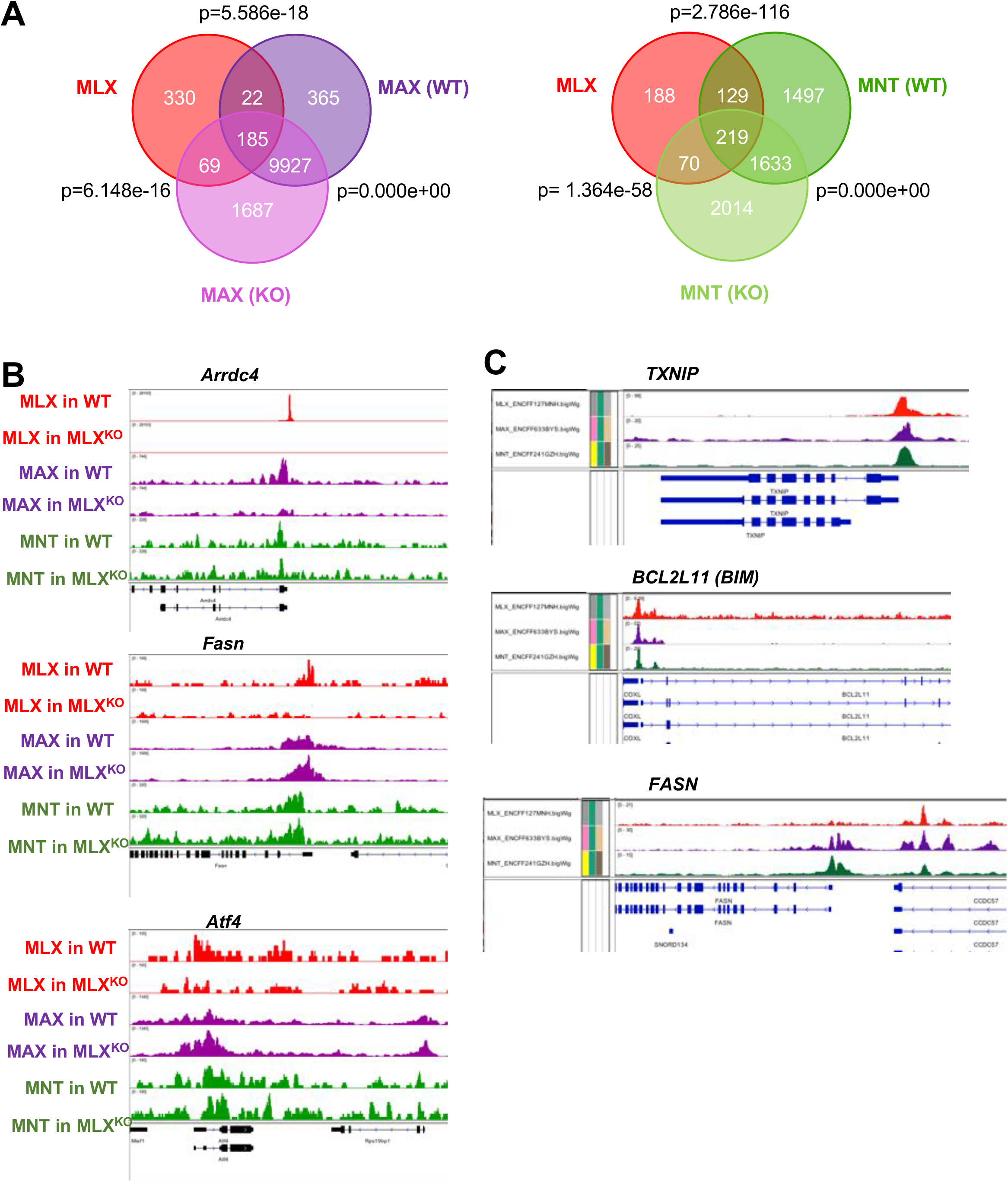
(Goes with Figure 6) MLX shares numerous transcriptional targets with MAX and MNT. **A**. Venn Diagram of the overlap of genes bound by MLX with MAX (Left), and MLX with MNT (Right), in both WT and MLX^KO^ 3T3 cells with p-value calculated from a hypergeometric test. **B**. IGV tracks for MLX, MAX and MNT on the *Arrdc4, Fasn and Atf4* promoters from WT and MLX^KO^ 3T3 cells. **C**. IGV tracks for MLX, MAX and MNT on the *TXNIP, BCL2L11 (BIM)* and *FASN* promoters from HepG2 cell ENCODE data.

**Supplemental Figure 7:**
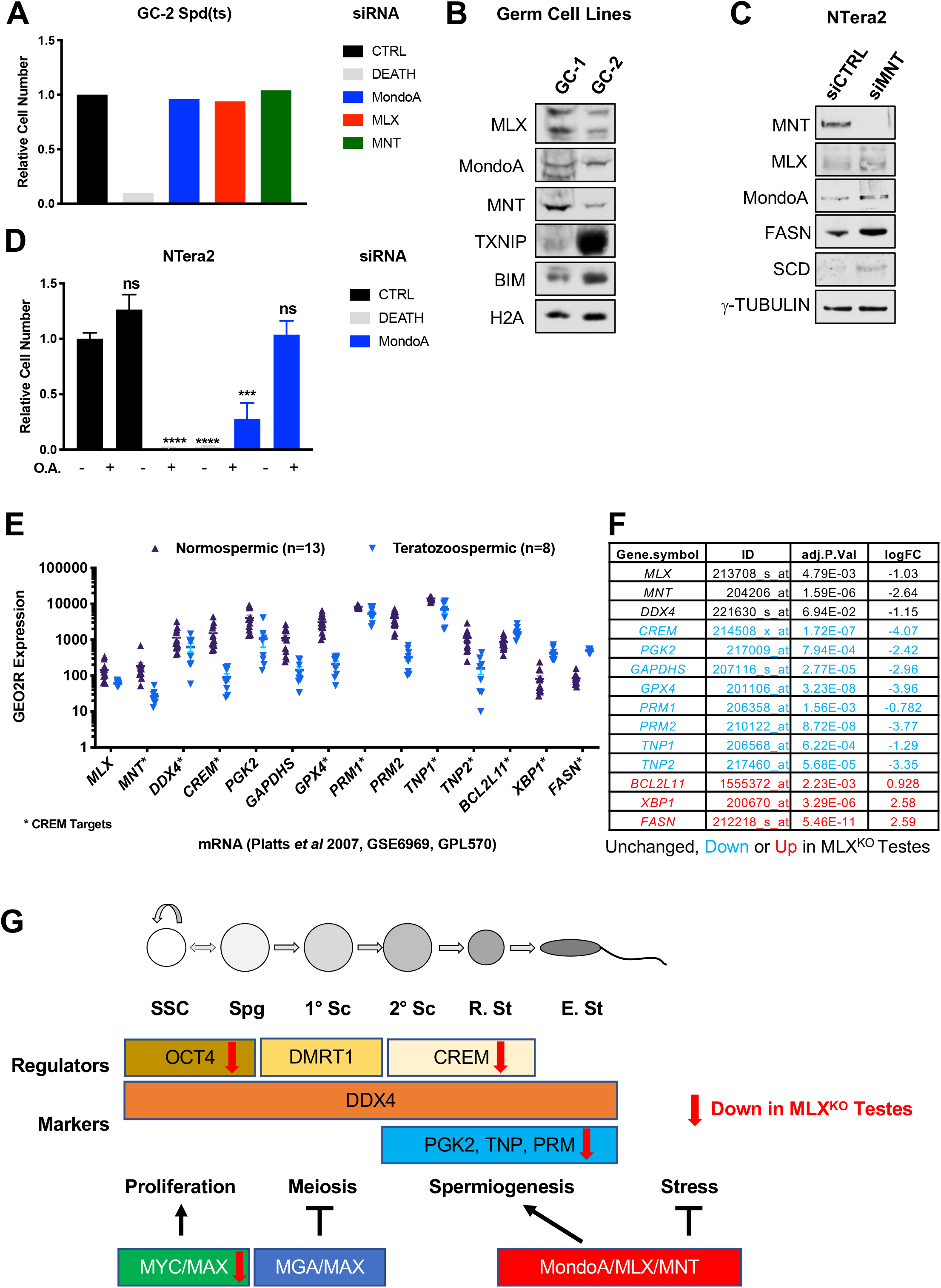
(Goes with Figure 7) MLX and MNT regulate metabolism and survival of male germ cell lines. **A**. Relative viable cell number of the GC-2 Spd(ts) cells after siRNA transfection with the indicated siRNA with siDEATH included as a control for siRNA transfection efficacy. **B**. Western blot analysis of GC-1-Spg and GC-2 Spd(ts) cells probed for the indicated proteins. **C**. Western blot analysis of NTera2 cells transfected with the indicated siRNA and probed for the indicated proteins. **D**. Relative viable cell number of the NTera2 cells after siRNA transfection with the indicated siRNA with siDEATH included as a control for siRNA transfection efficacy. Cells were cultured in the presence or absence of 35uM oleic acid O.A. (N=3 independent experiments, shown is the mean +/- SD). **E-F**. GEO2R analysis of GSE6969 from a published dataset of fertile (normospermic) versus infertile (teratozoospermic) men (Platts et al., 2007). LogFC and p-values are shown on the table to the right. **G**. Schematic of spermatogenesis indicating the stages likely to be influenced by MAX and MLX heterocomplexes.

